# Structural basis of ribosomal RNA transcription regulation

**DOI:** 10.1101/2020.06.05.136721

**Authors:** Yeonoh Shin, M. Zuhaib Qayyum, Danil Pupov, Daria Esyunina, Andrey Kulbachinskiy, Katsuhiko S. Murakami

## Abstract

Ribosomal RNA (rRNA) is the most highly expressed gene in rapidly growing bacteria and is drastically downregulated under stress conditions by the global transcriptional regulator DksA and the alarmone ppGpp. To reveal the mechanism of highly regulated rRNA transcription, we determined cryo-electron microscopy structures of the *Escherichia coli* RNA polymerase (RNAP) σ^70^ holoenzyme at different steps of rRNA promoter recognition with and without DksA/ppGpp. RNAP contacts the UP element of rRNA promoter using the dimerized α subunit carboxyl-terminal domain and scrunches the template DNA with the σfinger and β’lid to select a transcription start site favorable for rRNA expression. Promoter DNA binding to RNAP induces conformational change of the σ domain 2 that opens a gate for DNA loading and ejects σ_1.1_ from the RNAP cleft to facilitate open complex formation. DksA/ppGpp binding to RNAP also opens the DNA loading gate, but it is not coupled to σ_1.1_ ejection and impedes the open complex formation of the rRNA promoter due to its G+C rich discriminator sequence. Mutations in σ_1.1_ or the β’lid stabilize the RNAP and rRNA promoter complex and decrease its sensitivity to DksA/ppGpp. These results provide a molecular basis for exceptionally active rRNA transcription and for its vulnerability to DksA/ppGpp.

---

Bacteria sense the availability of nutrition and adjust ribosome biogenesis to optimize their growth. The rate of ribosome biogenesis is primarily determined by rRNA transcription ^1,2^, which constitutes as much as 70 % of total transcription and is initiated approximately every second from each of the seven operons (*rrn*A-E and *rrn*G-H) in *E. coli* during exponential growth ^3^. However, it is drastically repressed under stress conditions such as nutrient-starved stationary phase ^4^. rRNA expression is regulated at the initiation stage of RNA synthesis, including RNAP binding to promoter DNA, unwinding the DNA and escaping from the promoter.

The promoters (e.g., *rrnB*P1) for expressing rRNA operons are unique compared with other promoters, including 1) the A+T rich UP element located upstream of the −35 element (from −60 to −40); 2) the G+C rich discriminator sequence downstream of the −10 element (from −8 to −1); and 3) the transcription start site (TSS) located 9 bases downstream from the −10 element (**Fig. 1A, Supplemental Fig. 1A**). The UP element is recognized by the carboxyl-terminal domain of the α subunit (αCTD) and enhances rRNA transcription by more than 30-fold ^5^. The G+C rich discriminator and unusual TSS selection make the open complex (RPo) unstable, but this facilitates promoter escape by reducing the abortive RNA cycle prior to the productive RNA elongation stage^6^. These promoter elements play key roles in the wide range of rRNA transcription regulation between optimized and nonoptimized growth conditions.

**Figure 1.**
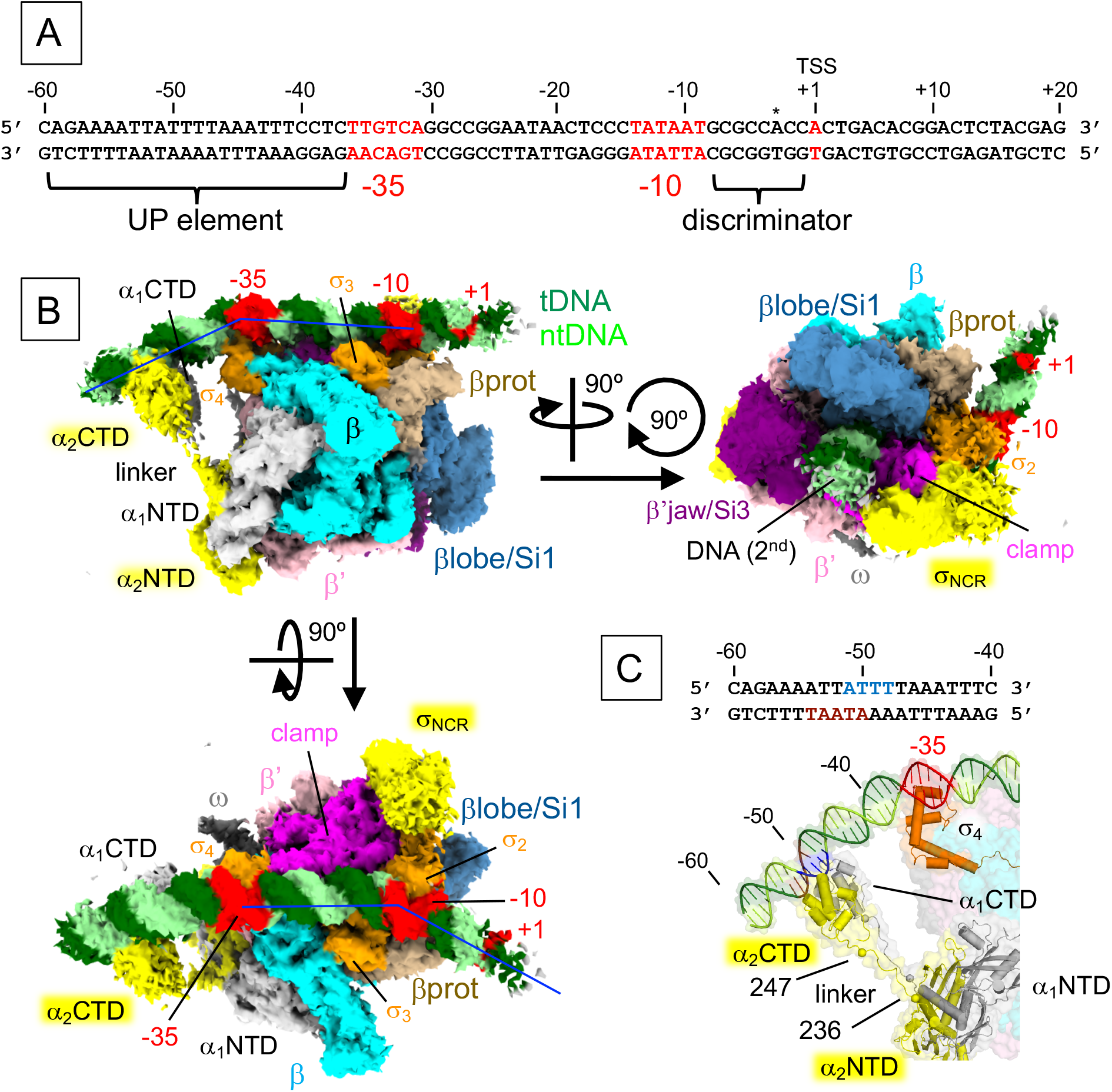
Cryo-EM structure of the RNAP - *rrnB* P1 closed complex (RPc). **A)** The sequence of the *E. coli rrnB*P1 promoter DNA used for cryo-EM. The UP element, −35 element, −10 element, transcription start site (TSS, +1) and discriminator sequence are indicated. An altered TSS from the nonscrunched open complex is indicated by an asterisk. **B)** Orthogonal views of the RPc cryo-EM density map. Subunits and domains of RNAP and DNA are colored and labeled (βprot, βprotrusion; tDNA, template DNA; ntDNA, nontemplate DNA). The density of downstream DNA beyond the +4 position is not traceable. Blue lines denote the direction of the DNA axis, with kinks at approximately −37 and −13. The second DNA at the RNAP cleft is indicated (DNA (2^nd^)). **C)** A magnified view showing the αCTDs and UP element interaction. The domains of α subunits, σ_4_ and DNA are depicted as ribbon models with a partially transparent surface. At the top, the sequence of the UP element is shown. The ntDNA (−51 to −48) and tDNA (−54 to −50) sequences binding α_1_CTD and α_2_CTD are highlighted in blue and brown, respectively.

rRNA transcription activity is regulated by the concentrations of two molecules - the initiating ribonucleotide (iNTP) (ATP in the case of *rrnB*P1) ^7^ and the bacterial alarmone ppGpp (guanosine tetraphosphate, aka “magic spot”), which is an allosteric effector of the RNAP-binding regulator DksA ^8–10^. The presence of high iNTP allows RNAP to break contacts with the promoter in the intrinsically unstable rRNA RPo, allowing escape to transcription elongation. However, the iNTP-limited condition shifts the equilibrium to favor early intermediates, including the closed complex (RPc), which is further enhanced by DksA/ppGpp binding to RNAP ^4^. The ppGpp concentration is increased under stress conditions, which enhances DksA-mediated rRNA transcription repression by stabilizing DksA in a functionally important binding mode.

The majority of bacterial RNAP-DNA complex structures determined by X-ray crystallography contain short promoter DNA fragments with a premelted transcription bubble that mimics RPo to maximize its stability ^11,12^. These studies explained the structural basis of promoter recognition and transcript initiation but left unexplored the interactions of RNAP with duplex DNA around the UP element (via αCTDs) and the contacts with the −10 element (via σ domain 2, residues 96-127 and 373-456 in σ^70^) in the RPc and the scrunched DNA bubble in the stressed RPo formed with the rRNA promoters.

Cryo-electron microscopy (cryo-EM) structures of the *E. coli* RNAP-*rpsT*P2 promoter complex with a DksA homolog TraR revealed the RPo formation pathway in the presence of TraR, with stepwise RNAP conformational changes ^13^. However, the *rpsT*P2 promoter for expressing ribosomal protein S20 is distinct from the *rrnB*P1 promoter in that it contains a G+C rich UP element and the TSS 7 bases downstream from the −10 element; therefore, it could not reveal the pathway for rRNA promoter complex formation and the mechanism of rRNA transcription regulation. Here, we used cryo-EM to visualize the RNAP and *rrnB*P1 complex in the RPc and RPo stages and two intermediates with DksA/ppGpp on the way to RPo formation.

## Cryo-EM structure of the RNAP and *rrnB*P1 promoter closed complex (RPc)

We preincubated RNAP with *rrnB*P1 promoter DNA (**Fig. 1A**) at 37 °C for 5 min prior to cryo-EM grid preparation. In a separate cryo-EM grid preparation, we also tested adding iNTPs (ATP and the nonhydrolyzable nucleotide CMPCPP) to stabilize the RNAP-*rrnB*P1 complex. In the course of cryo-EM data processing, 3D classification revealed 2 distinct structures based on the differences in the UP element (from −60 to −40), the downstream DNA (from −14 and +20) and the conformation of the σfactor (Methods, **SFigs. 2 and 3**), corresponding to the RPc and RPo. The first class represents the RNAP-*rrnB*P1 closed complex (RPc) with an overall resolution of 4.14 Å (**STable 1**). The cryo-EM density shows that RNAP binds the duplex DNA from −60 to +3 (**Fig. 1B, SFig. 4, SMovie 1**), but the density of downstream DNA beyond position +4 is not traceable. Instead, a second DNA binds to the RNAP cleft due to the ejection of σ_1.1_ from the RNAP cleft during RPc formation as described below.

**Figure 2.**
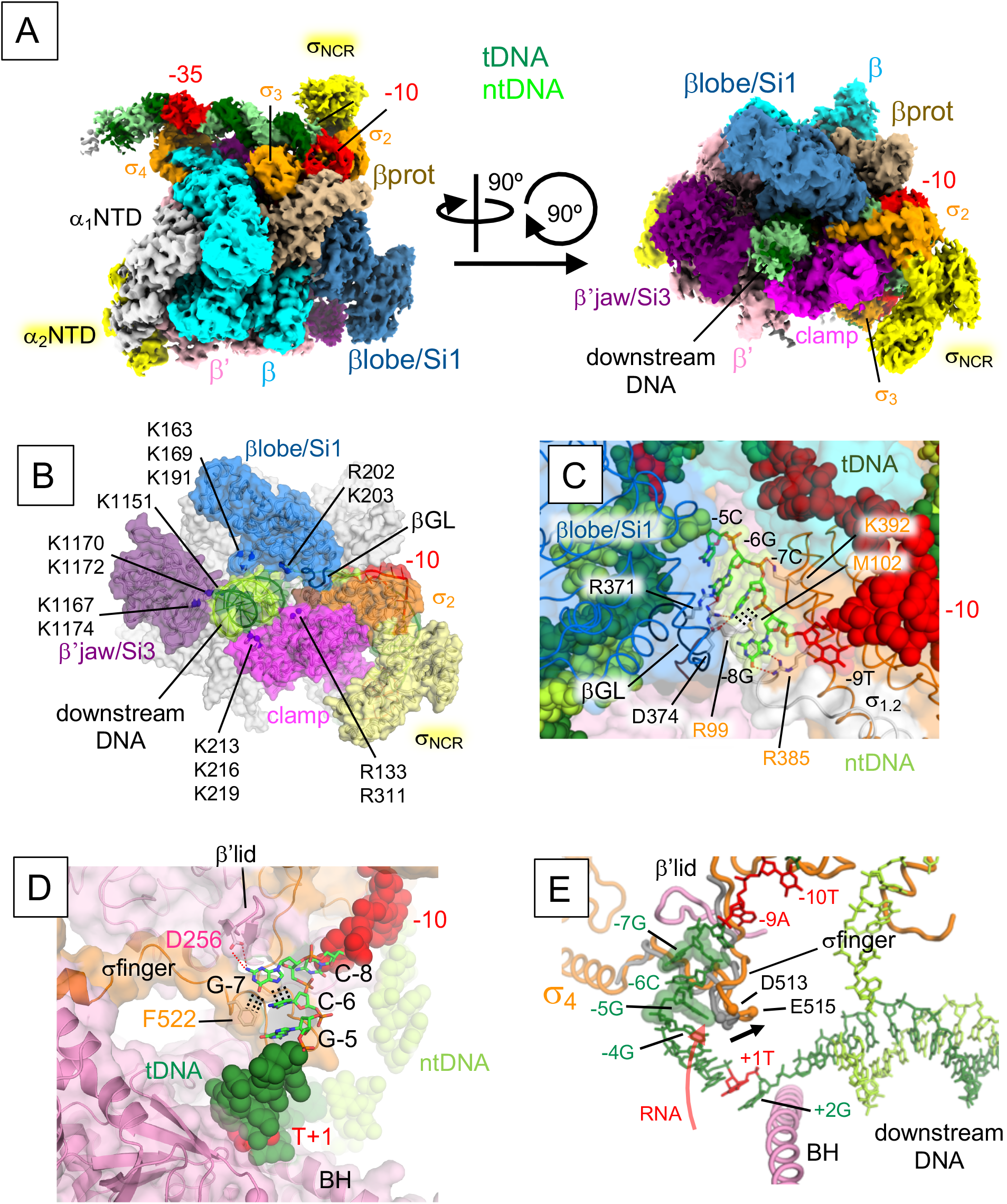
Cryo-EM structure of the RNAP – *rrnB* P1 open complex (RPo). **A)** Orthogonal views of the RPo cryo-EM density map. Subunits and domains of RNAP and DNA are colored and labeled the same as in Fig. 1B. **B)** The structure of the RPo, highlighting basic residues of the βlobe/Si1 (blue), β’jaw/Si3 (purple) and β’clamp (pink) interacting with downstream DNA (green) to stabilize the RPo. The structure is shown as a ribbon model with a transparent surface, and the basic residues are shown as spheres and labeled. **C)** Close-up view of RNAP (βGL, σ_1.1_ and σ_1.2_) and discriminator DNA (ntDNA) interaction. β and σ are depicted as ribbon models with transparent surfaces, and DNA is shown as CPK spheres. The G-8, C-7 and G-6 bases (stick model with transparent CPK spheres) that form salt bridges and Van del Waals interactions with residues from the βGL and σ_1.2_ (side chains shown as sticks; βGL R371 and D374; σ_1.2_ R99 and M102) are shown (depicted by red and black dashed lines). **D)** Close-up view of the RNAP (β’lid and σ finger) and discriminator DNA (tDNA) interaction. The G-7 base inserts into the pocket formed by the β’lid, σfinger and C-6 base. β’ and σ are depicted as ribbon models with transparent surfaces, and DNA is shown as a stick model and CPK representation. The residues forming salt bridges and Van del Waals interactions with the G-7 base are shown (depicted by red and black dashed lines). **E)** Comparison of the σfinger in RPo-*rrnB*P1 (this study, orange) and RPo-*rpsT*P2 ^13^ (gray). The RPo-*rrnB*P1 structure is depicted as cartoon (RNAP) and stick (DNA) models with the σfinger from RPo-*rpsT*P2 (gray). When the *rrnB*P1 tDNA scrunches at the −7G(t) position, −5G(t) is located below the σfinger (orange), which shifts the σfinger position compared to that in nonscrunched RPo. The σfinger dislocation (black arrow, 5 Å at E515) makes additional space for RNA extension (red arrow).

**Figure 3.**
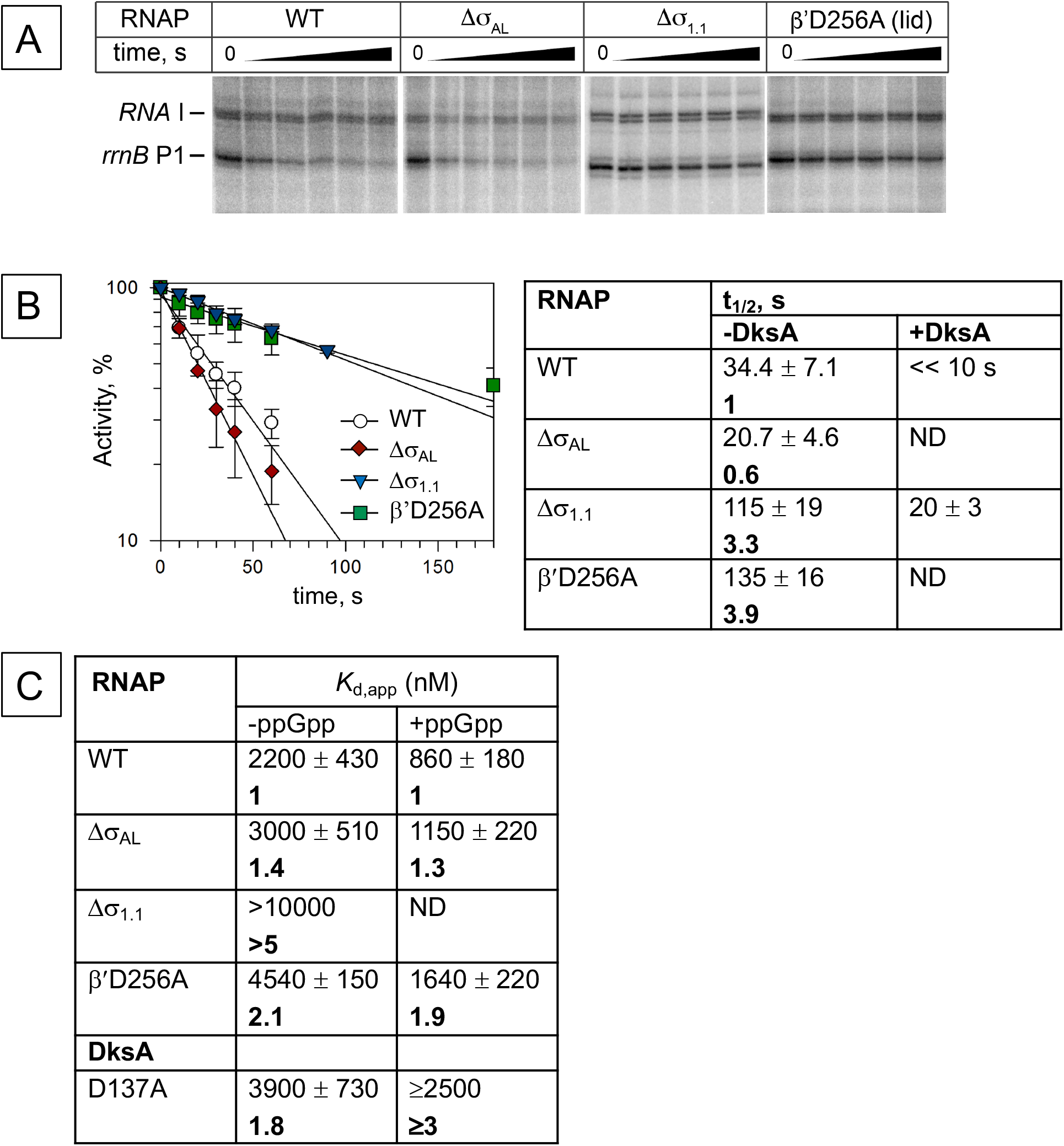
Stabilities of *rrnB* P1 promoter complexes formed by wild-type and mutant RNAPs and their sensitivities against DksA/ppGpp. **A**) Sensitivity of *rrnB*P1 promoter complexes to heparin. Preformed promoter complexes were incubated with heparin for the indicated time intervals, followed by the addition of NTPs and rifapentin. **B**) Kinetics of promoter complex dissociation for wild-type and mutant RNAPs. The half-lives of the promoter complexes for each RNAP are shown in the table to the right (mean values and standard deviations from three independent experiments). **C**) Apparent DksA affinities to wild-type and mutant RNAPs on the *rrnB*P1 promoter.

The cryo-EM density for both αCTDs (residues 248-329), the linkers (residues 236-247) connecting to αNTDs (residues 1-235), and the UP element DNA were traceable in the RPc, allowing us to investigate how each αCTD binds to the UP element unambiguously (**Fig. 1C, SMovie 1**). Two αCTDs form a head-to-tail dimer and bind DNA side-by-side in the middle of the UP element (−51 to −48 on nontemplate DNA (ntDNA) and −54 to −50 on template DNA (tDNA)), which is in good agreement with the DNA footprinting results ^14^. Although α subunits form a homodimer, two α subunits play different roles in RNAP, with one (α_1_) adjacent to the β subunit and the other (α_2_) adjacent to the β’. Compared to the α_2_CTD, the α_1_CTD is positioned proximally to the −35 element, which is consistent with the DNA cleavage by hydroxyl radicals from chelated Fe at each of the two αCTDs ^15^. The side chains of R265 and N294 from both αCTDs are inserted into the DNA minor groove, and basic residues (K291 and K298) are involved in salt bridges with the DNA phosphate backbone (**SFig. 5**). The linkers of both α subunits are fully extended, and slight DNA bending centered at the −37 position is required for the αCTDs binding to the UP element (**Fig. 1C**). Consistent with this observation, shortening of the linkers by only three amino acids reduces *rrnB*P1 transcription ^16^. Several studies have proposed that distant upstream DNA (near the −100 position) warps around RNAP on the RPo formation pathway, and the interaction of αCTDs and the UP element is one of the major driving forces for this DNA wrapping ^17,18^. However, αCTDs do not bend the DNA around its binding site in the RPc structure, indicating that these contacts may not contribute to the DNA wrapping by RNAP.

The RPc structure shows how σ_2_ binds the duplex form of the −10 element. The DNA encoding the −10 element is anchored by σ domain 2 and slightly bends around the upstream edge of the −10 element, allowing the downstream part beyond the −10 element to reach the other side of the RNAP cleft comprising the β protrusion domain (**Fig. 1B, SMovie 1**). The σregion 2.3 (σ_2.3_, residues 417-434) contacts the −10 element by fitting into the DNA major groove without any amino acid– DNA base interaction, indicating that σ_2.3_ recognizes the shape and/or curvature around the −10 element. This finding is in agreement with the previous proposal ^19^ that σ^70^ does not contact the - 10 element DNA bases when it is in duplex form.

## Cryo-EM structure of the RNAP and *rrnB*P1 promoter open complex (RPo)

The RNAP-*rrnB*P1 open complex (RPo) was determined with an overall resolution of 3.5 Å (**STable 1**). The cryo-EM density covers DNA from −44 to +20, including an open bubble from - 13 to +2 and the downstream DNA accommodated in the RNAP cleft (**Fig. 2A, SFig. 4, SMovie 2**). In contrast to the RPc, αCTDs and the UP element are disordered. Basic residues in the βlobe (K163, K169, K191, R202 and K203), β’jaw (K1151, K1167, K1170, K1172 and R1174) and β’clamp (R133, K213, K216, K219 and R311) participate in the interaction with downstream DNA to stabilize the RPo (**Fig. 2B**). The importance of these interactions in rRNA transcription regulation is supported by the isolation of Δ*dksA* suppressor mutations in these domains ^20,21^. The βgate loop (βGL, residues 368-378) in the βlobe domain contacts σregions 1.1 (σ_1.1_, residues 1-95) and 1.2 (σ_1.2_, residues 96-127) to enclose the RNAP cleft. The βGL, σ_1.2_ and σ_2.1_ (residues 373-396) contact the ntDNA strand of the discriminator from positions −8 to −6 (**Fig. 2C, SMovie 3**); consistently, the βGL deletion destabilizes the RPo and shifts the TSS to the −3A position ^22^. The RNAP and *rrnB*P1 complex starts RNA synthesis at the position 9 bp downstream from the - 10 element (+1A), which requires DNA scrunching in the open complex ^23^.

Nucleotide substitutions in several promoter positions, including the discriminator region, were shown to shift the TSS at *rrnB*P1 at position 6 bp downstream from the −10 element (−3A) and stabilize the complex, making it less sensitive to DksA/ppGpp. The RPo structure positioned +1A tDNA at the active site and revealed the mechanism of DNA scrunching, in which the G-7 base of tDNA fits into a pocket surrounded by the β’lid, σfinger (σ region 3.2) and C-6 base (**Fig. 2D, SMovie 3**). The importance of the G-7 base for the DNA scrunching is underscored by its conservation in all seven rRNA promoters in *E. coli* and some rRNA promoters in other proteobacteria (**SFig. 1**). Highly conserved D256 (β’lid) and F522 (σfinger) residues form a salt bridge and Van der Waals interaction with the G-7 base, respectively. Consistently, we found that an alanine substitution of residue D256 (β’lid) significantly stabilizes RNAP complexes with the *rrnB*P1 promoter (promoter complex half-life t_1/2_ of 135±16 s *vs.* 34±7 s for wild-type RNAP) (**Figs. 3A and B**). The σfinger deletion ^24^ or the G-7C substitution ^23^ was shown to shift the TSS to the −3A position, likely eliminating the open complex scrunching.

Open complex scrunching also facilitates the promoter escape of RNAP by reducing abortive RNA synthesis ^23^. Robust RNAP escape is required to initiate rRNA transcription approximately every second from each of the seven operons to meet the demand for ribosome synthesis in rapidly dividing *E. coli* ^3^. Compared with RPo-*rpsT*P2 containing nonscrunched tDNA ^13^, RPo-*rrnB*P1 shifts the σfinger ~5 Å away from tDNA, allowing accommodation of one additional base of RNA before its 5’-end reaches the σfinger (**Fig. 2E**). Since the σfinger is the major obstacle to promoter escape ^25–27^, the partially displaced σfinger in the rRNA promoter RPo due to open complex scrunching may decrease abortive RNA synthesis, promoting the robust expression of rRNA.

## Cryo-EM structures of the RNAP and *rrnB*P1 promoter complex with DksA/ppGpp (RP-DksA/ppGpp)

To reveal how DksA/ppGpp binding to RNAP downregulates rRNA transcription, we visualized the RNAP, *rrnB*P1 and DksA/ppGpp complex (RP-DksA/ppGpp) by cryo-EM (**STable 1, SFig. 6**). The classification of the cryo-EM data gave rise to two structures that differed mainly within the RNAP cleft; the first class shows the globular density corresponding to σ_1.1_ (class I, RP1-DksA/ppGpp), and the second class shows the right-handed helical density corresponding to the downstream DNA (class II, RP2-DksA/ppGpp) (**Figs. 4A and B, SMovie 4**). In addition, the positions of βlobe/Si1 are different in these classes (**Fig. 4C**).

**Figure 4.**
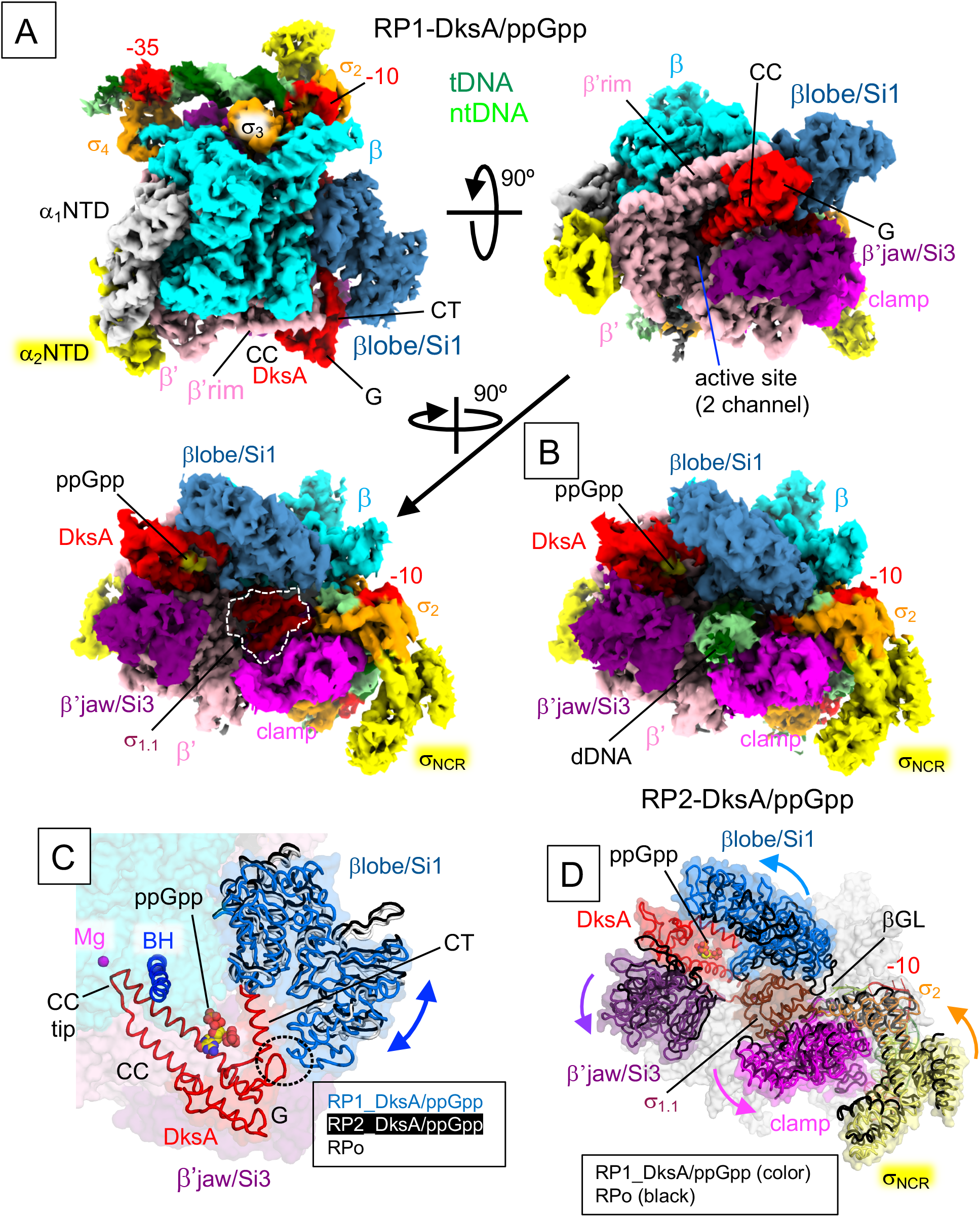
Cryo-EM structures of the RNAP - *rrnB* P1 complex with DksA/ppGpp (RP-DksA/ppGpp). **A)** Orthogonal views of the RP1-DksA/ppGpp cryo-EM density map. DNA, RNAP and DksA (G, G domain; CC, CC domain; CT, CT-helix) are indicated and colored. σ_1.1_ is highlighted by a white dash. **B)** The RP2-DksA/ppGpp cryo-EM density map. The downstream DNA is accommodated in the RNAP cleft. **C)** Close-up view of the βlobe/Si1 conformational changes upon DksA binding, σ_1.1_ ejection and downstream DNA binding. The structures of the βlobe/Si1 in RP1-DksA/ppGpp (light blue), RP2-DksA/ppGpp (white) and RPo (black) are depicted as ribbon models with transparent surfaces and ribbon models (DksA, BH: bridge helix) of RP1-DksA/ppGpp. The interaction between βlobe/Si1 and DksA in RP1-DksA/ppGpp is highlighted by a black dashed oval. **D**) Conformational changes in the RNAP mobile domains upon binding of DksA/ppGpp. The structures of the RNAP mobile domains in RP1-DksA/ppGpp (colored) and RPo (black) are depicted as ribbon models with transparent surfaces of RP1-DksA/ppGpp.

**Figure 5.**
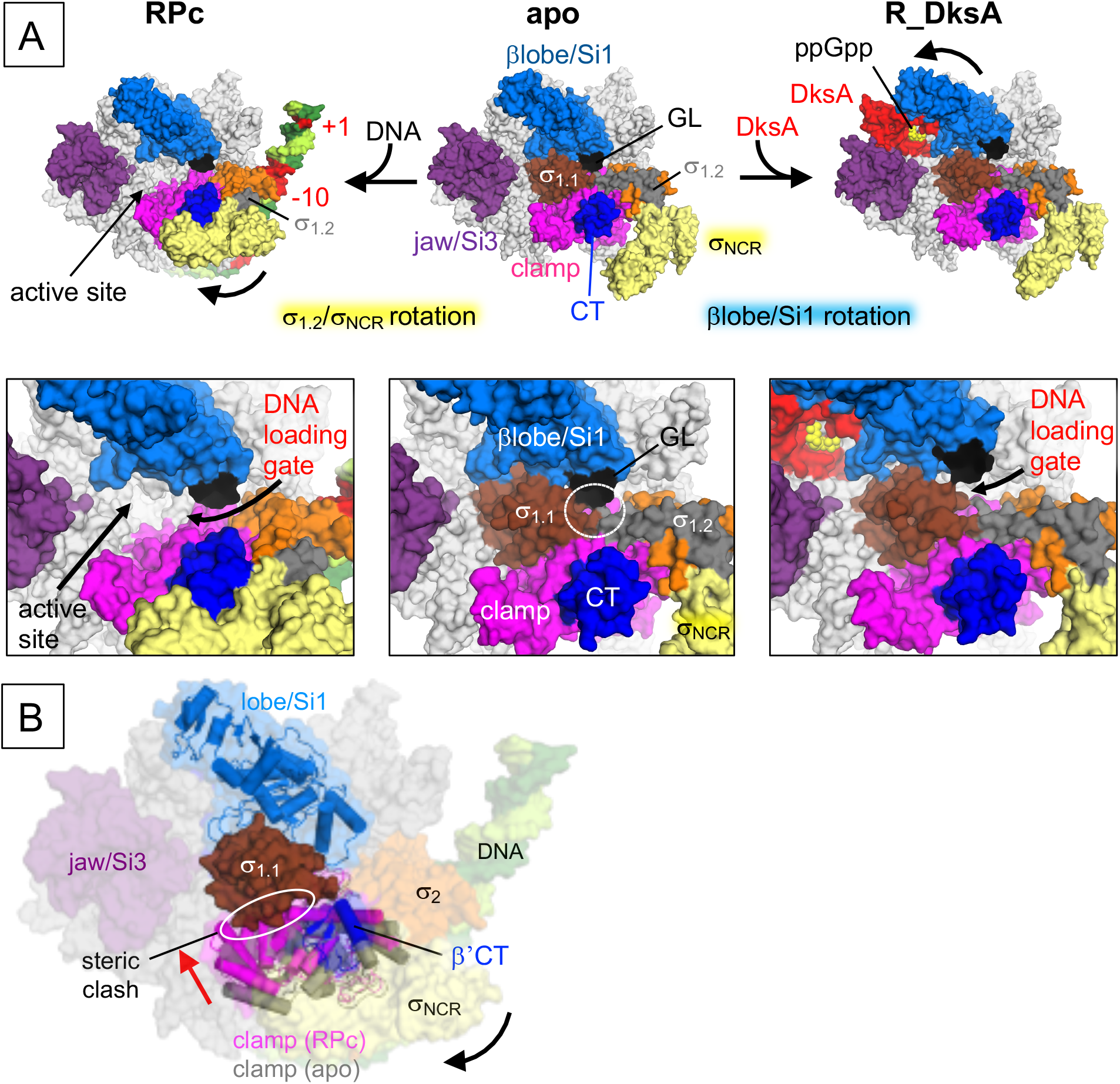
Opening the DNA loading gate by moving σ_1.2_/σ_NCR_ or βlobe/Si1 domain. **A)** Comparison of the σ_1.2_/σ_NCR_ and βlobe/Si1 conformations in apo-RNAP (middle), RPc (left) and R_DksA/ppGpp (right, DNA is removed from the RP1_DksA/ppGpp). RNAP (subunits and domains), DksA and DNA are indicated. Close-up views of the RNAP cleft are shown below. The DNA loading gate is closed in the apo-RNAP due to βgate loop (βGL) contacts σ_1.1_/σ_1.2_ (white dashed oval). The opening of the DNA loading gate in RPc and RP1_DksA/ppGpp is indicated by blue and red arrows, respectively. **B)** A proposed model of σ_1.1_ ejection in the RPc. The RPc is depicted as a transparent surface with cartoon models of the clamp (purple) and lobe/Si1 (blue). The clamp in an apo-form RNAP and the lobe/Si1 in RP1-DksA/ppGpp are colored gray and white, respectively. In the RPc, the σ_NCR_ rotation (back arrow) contacts the β’CT, resulting in clamp movement toward σ_1.1_ (red arrow) and a steric clash with σ_1.1_ (white oval).

Both classes show ppGpp binding at sites 1 and 2 and DksA binding at the RNAP secondary channel, as observed in a previous X-ray crystallography study ^28^. DksA binds RNAP with its globular domain (G domain, contacts with the β’rim helix), coiled-coil tip (CC tip, contacts with the active site), CC (contacts with the bridge helix, the trigger loop and linkers connecting to the β’Si3), and C-terminal α helix (CT-helix, contacts with the β lobe/SI1 domain) (**Fig. 4A, SMovie 4**). The CC of DksA prevents trigger helix formation and blocks NTP entry from the secondary channel, indicating that DksA must be displaced before RNAP can initiate RNA synthesis ^28,29^.

Both classes show the duplex DNA density from positions −42 to −14 (from the downstream edge of the UP element to the upstream edge of the −10 element) and also show the ssDNA density of the nontemplate strand of the −10 element (**SFig. 4**). RP1-DksA/ppGpp retains σ_1.1_ in the RNAP cleft, indicating that it represents an early stage intermediate during the RPc to RPo transition. While the transcription bubble is likely partially open in RP1-DksA/ppGpp, the density of ntDNA from −5 to +20 and of tDNA from −13 to +20 is not traceable. Analysis of RP1-DksA/ppGpp reveals a DksA/ppGpp-induced conformational change in βlobe/Si1, β’jaw/Si3 and β’clamp, opening the downstream DNA cleft in RNAP and likely reducing the stability of RPo (**Fig. 4D**). Furthermore, the conformational change in βlobe/Si1 establishes a new contact with the DksA CT-helix (**Fig. 4C, SMovie 4**); the deletion of βSi1 reduces the DksA affinity to RNAP and impairs its function ^30^. Alanine substitution of an aspartate residue in the CT-helix directly involved in this interaction (D137A) decreases *rrnB*P1 inhibition by DksA both in the absence and in the presence of ppGpp (**Fig. 3C**).

The RP2-DksA/ppGpp complex contains downstream DNA (from +3 to +20) within the RNAP cleft, but the density of the DNA bubble (from −8 to +2) is not traceable (**Fig. 4B, SFig. 4**), suggesting that it represents a late stage intermediate before forming the RPo. The σ_1.1_ density is not traceable due to its ejection from the RNAP cleft. The conformations of βlobe/Si1 and β’clamp are akin to the RPo conformation, and the CT-helix of DksA does not contact with the βlobe/Si1 (**Fig. 4C**). Therefore, the transition between the two complexes may reduce the DksA affinity to RNAP and trigger its dissociation, which is an obligatory process to initiate RNA synthesis ^28,29^.

## Conformational change in the σ domain 2 is coupled to σ_1.1_ ejection during RPc formation

The RPc structure revealed a significant conformational change in the σ domain 2 (from σregions 1.2-2.4 including σnonconserved region (σ_NCR_, residues 128-372)) comparison with the apo-form holoenzyme RNAP ^31^ or the RPo containing *rrnB*P1 (this study). Particularly, σ_1.2_/σ_NCR_ undergo a rigid rotation toward the clamp to establish contact with the β’clamp-toe (β’CT, residues 143 to 180) (**RPc, Fig. 5A**). Although this interaction was not observed in any previous structural study, it was predicted based on the biochemical/genetic analysis of RNAP promoter escape and early elongation pausing ^32^. It was shown that the interaction of the σ_NCR_ and β’CT is important for promoter escape and hinders early elongation pausing, and amino acid substitutions at the interface modulate both processes (**SFig. 7**).

The βGL contacts σ_1.1_ and σ_1.2_ to enclose the RNAP cleft in the apo-form RNAP, which prevents DNA loading (**apo, Fig. 5A**), but the same interaction in the RPo stabilizes the open complex bubble (**Fig. 2B**). In the case of RPc, the σ_1.2_/σ_NCR_ rotation disrupts the βGL and σcontact and widens the gap that allows discriminator DNA to enter the RNAP cleft for open complex bubble formation (**RPc, Fig. 5A**). Compared with the apo-form RNAP, the σ_NCR_ and β’CT interaction in the RPc closes the β’clamp, resulting in the ejection of σ_1.1_ from the RNAP cleft due to the steric clash between the β’clamp and σ_1.1_ (**Fig. 5B**).

σ_NCR_ contains a highly negatively charged region (acidic loop, residues 167-213) (**SFig. 8A**). Its conformation has not been determined due to its dynamic behavior, but since it is located near σ_2.3_, it seems to prevent nonspecific DNA binding to σ_2.3_ (**SFig. 8B**). We speculate that after RNAP recognizes the UP and −35 elements, loading of the −10 element DNA onto σ domain 2 triggers σ_NCR_ rotation due to charge-charge repulsion. After DNA unwinds around the −10 element, σ_NCR_ returns to its position, as seen in the RPo akin to the apo-form RNAP, and may enhance the electrostatic interaction between σ_2_ and −10 element DNA (**SFig. 8C**). Consistently, deletion of the acidic loop (Dσ_AL_) had a weak destabilizing effect on the *rrnB*P1-RNAP complex, without strong effects on DksA inhibition (**Fig. 3**).

DksA/ppGpp binding to RNAP also partially opens the DNA loading gate by moving the βlobe/Si1 away from σ_1.1_/σ_1.2_, but it is not coupled to the σ_1.1_ ejection from the RNAP cleft (**R_DksA, Fig. 5A**). Similarly, the structures of the RNAP-TraR complex and several RNAP-DNA complex intermediates prepared in the presence of TraR also showed the opening of the DNA loading gate by shifting the βlobe/Si1 position but did not show σ_1.2_/σ_NCR_ rotation or σ_1.1_ ejection from the RNAP cleft (**SFig. 9A**) ^13,31^.

To understand the role of σ_1.1_ in rRNA transcription, we characterized an RNAP derivative lacking σ_1.1_ (Δσ_1.1_-RNAP) in terms of its *rrnB*P1 transcription activity and sensitivity to DksA. Compared to the wild-type (WT) RNAP, Δσ_1.1_-RNAP increases *rrnB*P1 complex stability, both in the absence of DksA (increase t_1/2_ from 34 s to 115 s) and in its presence (increase t_1/2_ from ≪10 s to 20 s) (**Figs. 3A and B**), and it decreases sensitivity to DksA (**Fig. 3C**). The results indicate that σ_1.1_ plays an important role in rRNA transcription and its regulation by DksA/ppGpp.

## Mechanism of rRNA-specific transcription inhibition by DksA/ppGpp

Structural and biochemical studies of bacterial RNAP transcription suggest that the order of DNA loading around the TSS and DNA opening during promoter recognition may be interchangeable (i.e., DNA melts first outside RNAP (melt-load) or DNA melts after loading inside the RNAP cleft (load-melt)) depending on σ factors, promoters, transcription factors and conditions ^26,33^. By combining structural and biochemical data from this and previous studies, we propose two pathways of RPo formation (**Fig. 6, SMovie 5**). We hypothesize that RNAP uses alternative mechanisms of RPo formation, requiring opening of the DNA loading gate (disrupting the βGL contact to σ), σ_1.1_ ejection from the DNA binding channel, and unwinding of the −10 element plus discriminator DNA, depending on the absence or presence of DksA/ppGpp.

**Figure 6.**
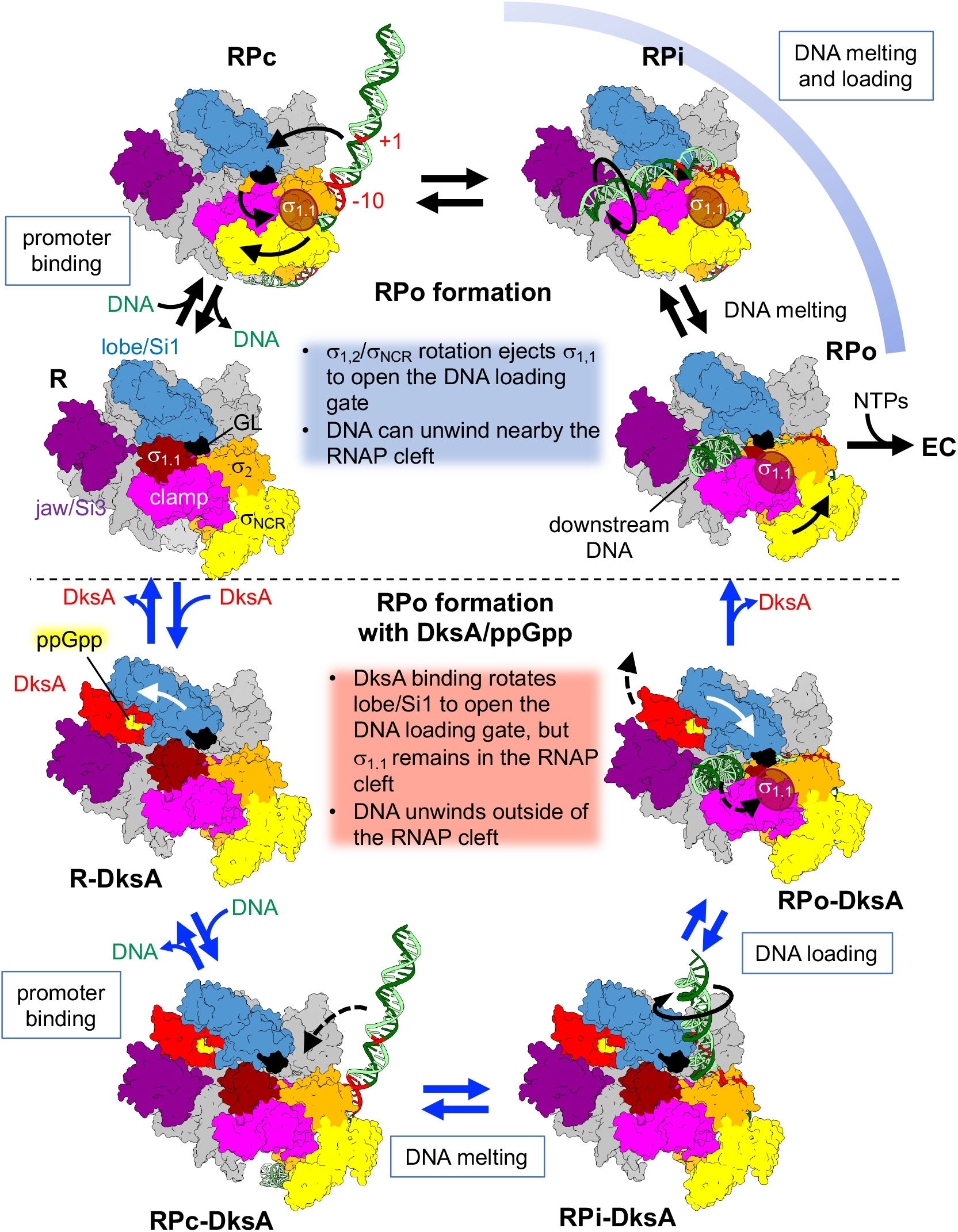
Alternative pathways for open promoter complex formation. Two distinct pathways are shown for open promoter complex formation without (top, blue caption) and with DksA/ppGpp (bottom, red caption). The RNAP holoenzyme (clamp, pink; σ_NCR_, yellow; σ_1.1_, brown; σ_2_, orange; βlobe/Si1, light blue; jaw/Si3, purple; rest of RNAP, gray), promoter DNA (tDNA, dark green; ntDNA, light green), DksA (red), and ppGpp (yellow) are shown. Only RPi is a hypothetical intermediate, but others (RPc, RPo, R-DksA, RPc-DksA, RPi_DksA and RPo-DksA) represent the structures determined in this and previous studies. EC, elongation complex.

Without DksA/ppGpp (**top, RPo formation**), free RNAP (**R**) binds promoter DNA (**RPc**), which opens the DNA loading gate by ejecting σ_1.1_ from the RNAP cleft, making RNAP competent for melting and loading discriminator DNA (**RPi**) into the RNAP cleft, which results in efficient RPo formation. The scrunched open complex (**RPo**) releases RNAP from the rRNA promoter rapidly to proceed with RNA synthesis (**EC**).

In the presence of DksA/ppGpp (**bottom, RPo formation with DksA/ppGpp**), DksA/ppGpp binding to RNAP rotates the βlobe/Si1 to DksA, which partially opens the DNA loading gate by disrupting the interaction between GL and σ(**R-DksA**). However, σ_1.1_ ejection is uncoupled from RPc formation (**RPc-DksA**), and σ_1.1_ remains inside the RNAP cleft until late stages of the open complex formation (**RPi-DksA**). This pathway favors the melt-load model for RPo formation (**RPo-DksA**), in which DNA is accommodated above the βlobe domain and unwinds outside the RNAP cleft (**SFig. 9B**) followed by single-stranded tDNA entry into the active site of RNAP ^13^. DNA unwinding outside the RNAP cleft is unfavorable in the DksA/ppGpp-free RNAP due to a steric clash of the discriminator DNA with the βlobe. The progression of DNA unwinding from the −10 element to the TSS is energetically less favorable for DksA/ppGpp-sensitive promoters (e.g., *rrnB*P1 and *rpsT*P2) containing the G+C rich discriminator than for less DksA/ppGpp-sensitive promoters (e.g., T7A1 and RNA1) containing an A+T rich discriminator (**SFig. 10**). *E. coli* promoters that are sensitive to DksA/ppGpp contain G+C rich discriminators ^34^. Replacing the A+T rich discriminator of the *uspA* promoter, which is positively regulated by DksA/ppGpp, with the one from the *rrnB*P1 promoter makes the *uspA* hybrid promoter sensitive to DksA/ppGpp ^35^, indicating that discriminator sequences play an important role in responding to DksA/ppGpp. Although DksA could inhibit transcription regardless of the promoter bound to RNAP, by inhibiting NTP entry and folding trigger helix, stable promoter complex formation decreases DksA binding to RNAP, thus relieving the inhibition ^28,29^. The completion of discriminator DNA loading into the RNAP cleft can likely occur not only in DksA/ppGpp-insensitive promoters but also in a fraction of the rRNA promoter complexes to maintain a basal level of rRNA expression under stress growth conditions. This likely pushes the βlobe/Si1 away from the CT-helix of DksA (**RPo-DksA**), allowing rapid dissociation of DksA from the RNAP secondary channel (**RPo**) followed by initiate transcription (**EC**).

In this study, we revealed two alternative pathways for opening the gate of the DNA binding channel depending on the absence or presence of DksA/ppGpp (**Fig. 6, SMovie 5**) and shed light on the functions of σ_1.1_, σ_1.2,_ σ_NCR_ and βlobe/Si1 domains to explain how DksA/ppGpp specifically inhibits rRNA transcription. Intriguingly, DksA/ppGpp is able to activate transcription at some σ^70^-promoters ^36^ and promoters recognized by alternative σ factors, including σ^S 37^ and σ^E 38^. Neither σ^S^ nor σ^E^ contains σ_1.1_ or σ_NCR_, and the σ^S^ and σ^E^ holoenzymes use the βGL to close the DNA loading gate ^39,40^. DNA binding to the σ domain 2 of σ^S^ or σ^E^ cannot facilitate opening of the DNA loading gate, as described in the case of the σ^70^ holoenzyme (**Fig. 6 top**). However, DksA/ppGpp binding followed by movement of the βlobe/Si1 domain could still open the DNA loading gate of these RNAP holoenzymes as described above (**Fig. 6 bottom**), possibly explaining the stimulatory effects of DksA/ppGpp on transcription from some σ^S^- and σ^E^-dependent promoters. Further structural analyses of the σ^70^, σ^S^ and σ^E^ RNAP promoter complex with DksA/ppGpp, together with biochemical characterization of RNAP and promoter functions, will be needed to complete our understanding of DksA/ppGpp-dependent transcription regulation.

## Supporting information

SMovie1

SMovie2

SMovie3

SMovie4

SMovie5

## REFERENCES

Uncategorized References

## METHODS

### Purification of *E. coli* WT RNAP, RNAP derivatives, and DksA

*E. coli* σ^70^ RNAP holoenzyme and DksA were purified as described previously ^28,41^. *E. coli* RNAP derivatives were purified by the same method as WT RNAP.

### Preparation of *rrnB*P1 DNA

The *rrnB*P1 promoter DNA was synthesized (IDT) according to the native *rrnB*P1 sequence and annealed in a 40 μL reaction mixture containing 10 mM Tris-HCl (pH 8.0), 50 mM NaCl, and 1 mM EDTA to a final concentration of 0.5 mM. The solution was heated at 95 °C for 10 min, and then the temperature was gradually decreased to 22 °C. The sequence of the nontemplate strand is 5’-CAGAAAATTATTTTAAATTTCCTCTTGTCAGGCCGGAATAACTCCCTATAATGCGCC ACC<u>A</u>CTGACACGGACTCTACGAG-3’. The transcription start site is underlined, and the template sequence is 5’-CTCGTAGAGTCCGTGTCAGTGGTGGCGCATTATAGGGAGTTATTCCGGCCTGACAAGAGGAAATTTAAAATAATTTTCTG −3’.

### Cryo-EM sample preparation

To prepare the RNAP and *rrnB*P1 promoter complex, *E. coli* σ^70^ RNAP (20 μM) and *rrnB*P1 promoter DNA (40 μM) were preincubated for 5 min at 37 °C in buffer (10 mM HEPES, pH 8.0, 50 mM NaCl, 0.1 mM EDTA, 5 mM DTT and 5 mM MgCl_2_) followed by the addition of ATP and a nonhydrolyzable CMPCPP (cytidine-5’-[(α,β)-methyleno]triphosphate, Jena Bioscience) (2 mM each). After mixing, the reaction was further incubated for 5 mins at 37 °C.

To prepare RP-DksA/ppGpp, *E. coli* σ^70^ RNAP (20 μM) was preincubated with a 5-fold molar excess of DksA (100 μM) and ppGpp (2 mM) for 5 min at 37 °C in buffer (10 mM HEPES, pH 8.0, 50 mM NaCl, 0.1 mM EDTA, 5 mM DTT and 5 mM MgCl_2_). The *rrnB*P1 promoter DNA (40 μM) was added to the reaction and further incubated for 5 mins at 37 °C. Before freezing the grids, 8 mM CHAPSO (Hampton research) was added to the reaction. A 3.5 μL sample was applied to a glow-discharged C-Flat Holey Carbon grid (Cu 2/1, 400 mesh), blotted and plunge-frozen in liquid ethane using a Vitrobot Mark IV (FEI, USA) with 95% humidity at 4 °C.

### Cryo-EM data acquisition

The grid was imaged using a 300 keV Titan Krios (Thermo Fisher) microscope equipped with a K3 direct electron detector (Gatan) and controlled by the Latitude S (Gatan, Inc.) software at the National Cancer Institute’s Cryo-EM Facility at Frederick. The defocus range was 1.0 – 3.0 μm, and the magnification was 81,000X in electron counting mode (pixel size = 1.08 Ǻ/pixel). Forty frames per movie were collected with a dose of 1.125 e^−^/Å^2^/frame, giving a total dose of 45 e^−^/Å^2^.

### Cryo-EM data processing

The RNAP-*rrnB*P1 complex with ATP/CMPCPP data was processed using Relion3.0.8 ^42^. A total of 8,315 movies were collected, aligned and dose weighted using MotionCor2 ^43^. CTF fitting was performed with Gctf ^44^. Initially, approximately 1,000 particles were manually picked to generate particle templates followed by automated picking, resulting in a total of 1,449,010 particles subjected to 2D classification. From the 2D classes, 1,442,810 particles were chosen for the 3D classification to 4 classes. Poorly populated classes were removed, resulting in datasets of 541,257 (37%) particles for the first class (RPc) and 464,512 (32%) particles for the second class. The first class was further 3D classified without alignments twice to further clean the data, resulting in datasets of 67,187 particles. The particles were refined and postprocessed to generate the density map at 4.14 Å resolution. The resolution of the density map of the second class was 3.53 Å.

The RNAP-*rrnB*P1 complex data were processed using Relion3.0.8. A total of 4748 movies were collected, aligned and dose weighted using MotionCor2. CTF fitting was performed with Gctf. Approximately 1,000 particles were manually picked to generate particle templates followed by automated picking, resulting in a total of 563,500 particles. Particles were 2D classified, and 561,753 particles were chosen for the 3D classification. Of the four 3D classes, class 1 (RPo) was the most populated class (349,752 particles, 62%) and was autorefined. The map was postprocessed to give a structure of RPo at 3.53 Å.

The RP-DksA/ppGpp complex data were processed using cryoSPARC V2.9.0 ^45^. A total of 4,926 movies were collected, and the movies were aligned, and dose weighted using Patch-motion correction. CTF fitting was performed with Patch-CTF estimation. Initially, approximately 1,000 particles were manually picked to generate particle templates followed by automated picking, resulting in a total of 418,049 particles subjected to 2D classification. After two rounds of 2D classification to remove junk particles, 361,048 particles were used to generate two *ab initio* models. Junk particles were removed, resulting in a dataset of 275,629 particles chosen for the 3D classification (heterogenous refinement). Poorly populated classes were removed, resulting in a dataset of 49,995 particles to generate the density map at 3.62 Å resolution for the first class (RP1-DksA/ppGpp) and a dataset of 79,275 particles to generate the density map at 3.58 Å resolution for the second class (RP2-DksA/ppGpp). The particles were 3D autorefined without the mask and postprocessed (homogenous refinement).

### Structure refinement

To refine the closed and open complex structures, the *E. coli* RNAP holoenzyme crystal structure (PDB: 4YG2) was manually fit into the cryo-EM density map using Chimera ^46^ and real-space refined using Phenix ^47^. In the real-space refinement, the domains of RNAP were rigid-body refined and then subsequently refined with secondary structure, Ramachandran, rotamer and reference model restraints. To refine the structures of RP1-DksA/ppGpp and RP2-DksA/ppGpp, *E. coli* RNAP and DksA/ppGpp complex crystal structures (PDB: 5VSW) were manually fit into the cryo-EM density map using Chimera. DNA was manually built by using Coot ^48^. The structure was refined by the same method as the closed and open complex structures.

### Preparation of RNAP and transcription factors for *in vitro* transcription

Mutant variants of RNAP, σ^70^ and DksA were obtained by site-directed mutagenesis. The D256A substitution in the β’ subunit was obtained in pVS10 encoding all RNAP subunits, with the *rpoC* gene containing a C-terminal His_6_-tag ^49^. The σ^70^ and DksA variants containing an N-terminal His_6_-tag were cloned into pET28. To obtain σΔ_1.1_, the 5’-terminal part of the *rpoD* gene encoding residues 2-94 was deleted. To obtain σΔ_AL_, codons 168-212 were replaced with three glycine codons. All proteins were expressed in *E. coli* BL21(DE3). The wild-type and mutant core RNAPs were purified using Polymin P precipitation followed by heparin (HiTrap Heparin column), Ni-affinity (HisTrap HP column) and anion exchange (MonoQ column) chromatography steps (all columns from GE Healthcare) as described previously ^49^. The wild-type and mutant σ^70^ factors were purified from inclusion bodies with subsequent renaturation and Ni-affinity chromatography as previously described ^27^. The σΔ_1.1_ protein was subjected to thrombin protease (GE Healthcare) treatment in PBS buffer (10 hours of incubation at 4 °C with 10 units of protease per mg of protein), followed by incubation with Ni-NTA agarose (GE Healthcare) to remove the His-tag and His-tagged thrombin. To purify DksA, a bacterial pellet from 0.5 liters of cell culture was resuspended in 25 ml of lysis buffer (50 mM Tris-HCl, pH 7.9, 250 mM NaCl, 10 mM EDTA, 0.5 mM phenylmethylsulfonyl fluoride, 1 mM 2-mercaptoethanol, 0.1 mM ZnCl_2_) and lysed using a French press. The supernatant obtained after centrifugation was loaded onto a 5-ml HiTrap chelating column (GE Healthcare) charged with Ni^2+^ and equilibrated with loading buffer (10 mM Tris-HCl, pH 7.9, 500 mM NaCl, 0.5 mM 2-mercaptoethanol, 0.1 mM ZnCl_2_). The column was washed with the same buffer containing 60 mM imidazole, and DksA was eluted with buffer containing 300 mM imidazole and dialyzed overnight against 50 mM Tris-HCl, 300 mM NaCl, 1 mM DTT, and 0.1 mM ZnCl_2_. Glycerol was added to 50%, and aliquots were stored at −70 °C.

### Transcription *in vitro*

Analysis of transcription *in vitro* was performed using a supercoiled pTZ19 template containing *rrnB*P1 cloned 88 nt upstream of a *his* terminator (Pupov et al., 2018); the second transcript monitored in the assays was 110 nt RNA-I encoded by the *ori* region of the plasmid. For measurements of promoter complex stabilities, promoter complexes were prepared by mixing core RNAP (100 nM final concentration) with wild-type or mutant σ^70^ factors (250 nM) in transcription buffer (40 mM Tris-HCl, pH 7.9, 10 mM MgCl_2_, 40 mM KCl) and supercoiled plasmid DNA (10 nM), followed by incubation for 7 min at 37 °C. DksA and ppGpp were added at 2 μM and 200 μM, respectively, when indicated. An upstream fork-junction competitor DNA was added (template strand 5’-ACGAGCCGGAAGCAT, nontemplate strand 5’-ATGCTTCCGGCTCGT<u>ATAA</u>TGTGTGGAA; the −10 sequence is underlined) to 2 μM, and the samples were incubated at 37 °C for the indicated time intervals. NTP substrates were added to final concentrations of 200 μM ATP, CTP, GTP, and 10 μM UTP, with the addition of α-[^32^P]-UTP together with rifapentin (5 μg/ml) to prevent reinitiation. The reactions were stopped after 5 min with 8 M urea and 20 mM EDTA, and RNA products were separated by 15% denaturing PAGE, followed by phosphor imaging. To calculate the observed half-life times for promoter complex dissociation (*t*_1/2_), the data were fitted to the one-exponential equation *A* = *A*_0_ × exp(–t × *k*_obs_), where A is the RNAP activity at a given time point after competitor addition, A_0_ is the activity measured in the absence of the competitor, *k*_obs_ is the observed rate constant, and *t*_1/2_ = ln2/*k*_obs_.

For measurements of apparent DksA affinities, promoter complexes were prepared in the same way with 50 nM core RNAP, 250 nM σ^70^ (250 nM) and 2 nM supercoiled plasmid DNA in transcription buffer containing 100 μg/ml BSA for 7 min at 37 °C, followed by the addition of DksA (from 10 nM to 10 μM), either in the absence or in the presence of ppGpp (100 μM). Transcription was performed for 15 min at 37 °C with 200 μM ATP, CTP, GTP, and 10 μM UTP (plus α-[^32^P]-UTP), and RNA products were analyzed as described above. The apparent dissociation constant values (*K*_d,app_) were calculated from the hyperbolic equation: A = A_max_ × (1 – [DksA]/(*K*_d,app_ + [DksA])), where A is the observed RNAP activity and A_max_ is the RNAP activity measured in the absence of DksA.

### DNA duplex free energy calculation

DNA duplex free energies were analyzed based on nearest-neighbor thermodynamics ^50,51^. Briefly, the Python script was written to read a sequence from a text file, calculate the DNA duplex free energy of dinucleotides, sum these values over an 8-base window and report these sums for the first base of the central nucleotide of the window (e.g., sum for the first window base 1-8 will be reported for base 4).

## Data Availability

The cryo-EM density maps have been deposited in EMDataBank under accession codes EMDB: EMD-21879 (RPc), EMD-21880 (RPo), EMD-21881 (RP1-DksA/ppGpp), and EMD-21883 (RP2-DksA/ppGpp). Atomic coordinates for the reported cryo-EM structures have been deposited with the Protein Data Bank under accession numbers 6WR6, 6WR8, 6WRD and 6WRG.

## ACKNOWLEDGMENTS

We thank Carol Bator at the Penn State Huck Institute Cryo-EM facility for supporting the cryo-EM data collections. We thank Christian Dienemann at the Max Planck Institute for providing the DNA duplex free energy calculation methodology. This research was, in part, supported by the National Cancer Institute’s National Cryo-EM Facility at the Frederick National Laboratory for Cancer Research under contract HSSN261200800001E. This work was supported by NIH grants (R01 GM087350 and R35 GM131860 to K.S.M.) and Russian Science Foundation (19-14-00359) and Russian Foundation for Basic Research (18-34-20095) to A.K.

## AUTHOR CONTRIBUTIONS

Y.S. prepared samples and cryo-EM grids. Y.S. and M.Z.Q. collected and processed cryo-EM data. K.S.M. built, refined and validated the structures. D.P. and D.E. performed biochemical assays under supervision of A.K. K.S.M. designed and supervised research. All authors participated in the interpretation of the results and in writing the manuscript.

## Correspondence and requests for materials

should be addressed to K.S.M. or A.K.

