## Supplemental info for "Structural basis of ribosomal RNA transcription regulation"

### EXTENDED DATA

#### STable 1. Cryo-EM data collection, refinement and validation statistics

| | RPc<br>(EMD-21879)<br>(6WR6) | RPo<br>(EMD-21880)<br>(6WR8) | RP1_DksA/ppGpp<br>(EMD-21881)<br>(6WRD) | RP2_DksA/ppGpp<br>(EMD-21883)<br>(6WRG) |
|---|---|---|---|---|
| **Data collection and processing** | | | | |
| Magnification | 81,000 | 81,000 | 81,000 | 81,000 |
| Voltage (kV) | 300 | 300 | 300 | 300 |
| Electron exposure ($e^-$/Å$^2$) | 45 | 45 | 45 | 45 |
| Defocus range (μm) | 1.0-2.5 | 1.0-2.5 | 1.0-2.5 | 1.0-2.5 |
| Pixel size (Å) | 1.08 | 1.08 | 1.08 | 1.08 |
| Symmetry imposed | C1 | C1 | C1 | C1 |
| Initial particle images (no.) | 1,442,810 | 561,753 | 275,629 | 275,629 |
| Final particle images (no.) | 67,187 | 349,752 | 49,995 | 79,275 |
| Map resolution (Å)<br>FSC threshold | 4.14<br>0.143 | 3.53<br>0.143 | 3.62<br>0.143 | 3.58<br>0.143 |
| Map resolution range (Å) | 3.7-10.0 | 3.8-11.0 | 3.2-7.8 | 3.1-8.0 |
| **Refinement** | | | | |
| Initial model used (PDB code) | 4YG2 | 4YG2 | 5VSW | 5VSW |
| Model resolution (Å)<br>FSC threshold | 4.1<br>0.143 | 3.5<br>0.143 | 3.6<br>0.143 | 3.6<br>0.143 |
| Map sharpening *B* factor (Å$^2$) | -110 | -125 | -75 | -80 |
| *Model composition*<br>Non-hydrogen atoms<br>Protein residues<br>Ligands | <br>33,509<br>3,830<br>Zn:2, Mg:1,<br>1N7:2, POP:1 | <br>31,608<br>3,688<br>Zn:2, Mg:1,<br>1N7:2 | <br>32,349<br>3936<br>G4P:2, Zn:3, Mg:1,<br>1N7:4 | <br>32,527<br>3850<br>G4P:2, Zn:3, Mg:1,<br>1N7:4 |
| *B factors (Å$^2$)*<br>Protein<br>Ligand | <br>81.31<br>60.80 | <br>77.99<br>72.16 | <br>98.22<br>99.83 | <br>105.05<br>104.34 |
| *R.m.s. deviations*<br>Bond lengths (Å)<br>Bond angles (°) | <br>0.011<br>1.166 | <br>0.009<br>0.900 | <br>0.009<br>1.011 | <br>0.012<br>1.192 |
| *Validation*<br>MolProbity score<br>Clash score<br>Poor rotamers (%) | <br>2.81<br>61.99<br>0.31 | <br>3.40<br>54.85<br>7.91 | <br>2.71<br>48.09<br>0.27 | <br>2.74<br>52.18<br>0.37 |
| *Ramachandran plot*<br>Favored (%)<br>Allowed (%)<br>Disallowed (%) | <br>89.97<br>9.27<br>0.76 | <br>91.38<br>8.02<br>0.60 | <br>89.84<br>9.11<br>1.05 | <br>90.11<br>9.11<br>0.78 |

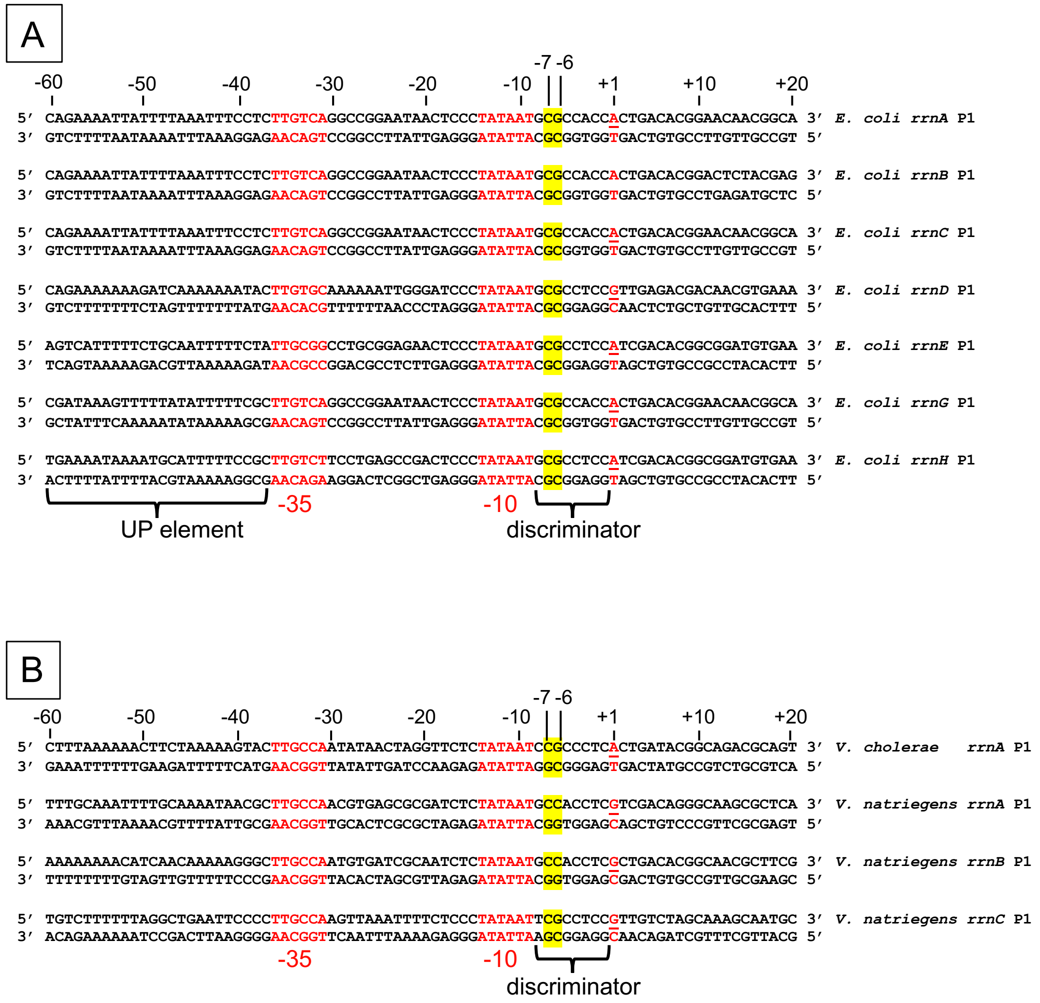

**SFigure 1.** ***rrnA-H*** **and other promoter sequences.**

**A**) Sequences of the *E. coli rrnA*P1 – *rrnH*P1 promoter DNAs [1,2]. The -35 element, -10 element and transcription start site (TSS, +1) are indicated in red, and the -7 and -6 bases are highlighted in yellow.

**B**) Representative rRNA promoter sequences from *Vibrio cholerae* and *Vibrio natriegens* (γ-proteobacteria) [3]. The -35 element, -10 element and TSS are indicated in red, and the -7 and -6 bases are highlighted in yellow.

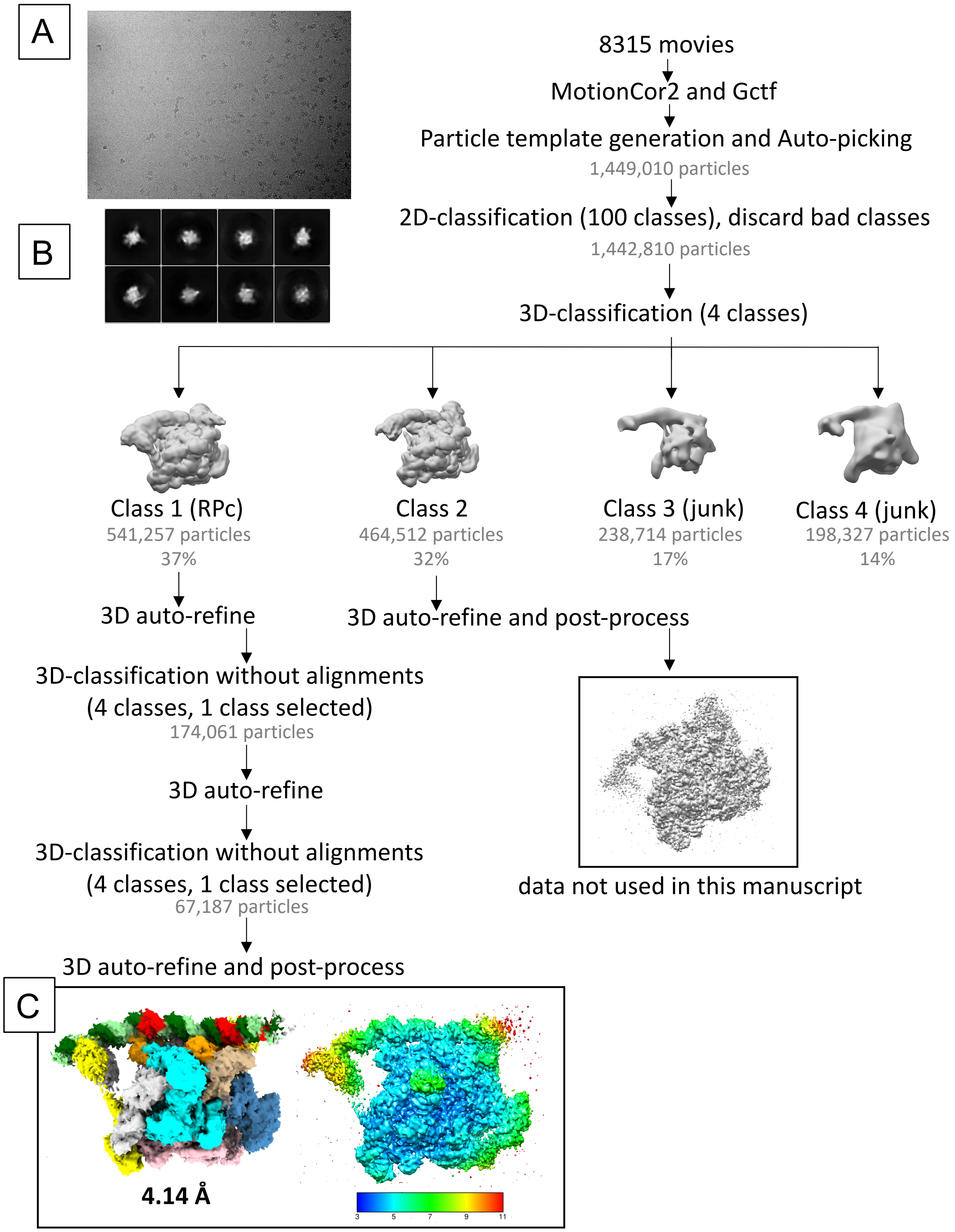

**SFigure 2. Cryo-EM processing pipeline for RNAP-*rrnB*P1 complex with ATP/CMPCPP.**

**A**) A representative micrograph used for data processing.

**B**) Selected representative 2D classes from 2D classification.

**C**) RPc. The cryo-EM density map is colored according to Fig. 1B. The right view is the same as on the left but colored by local resolution.

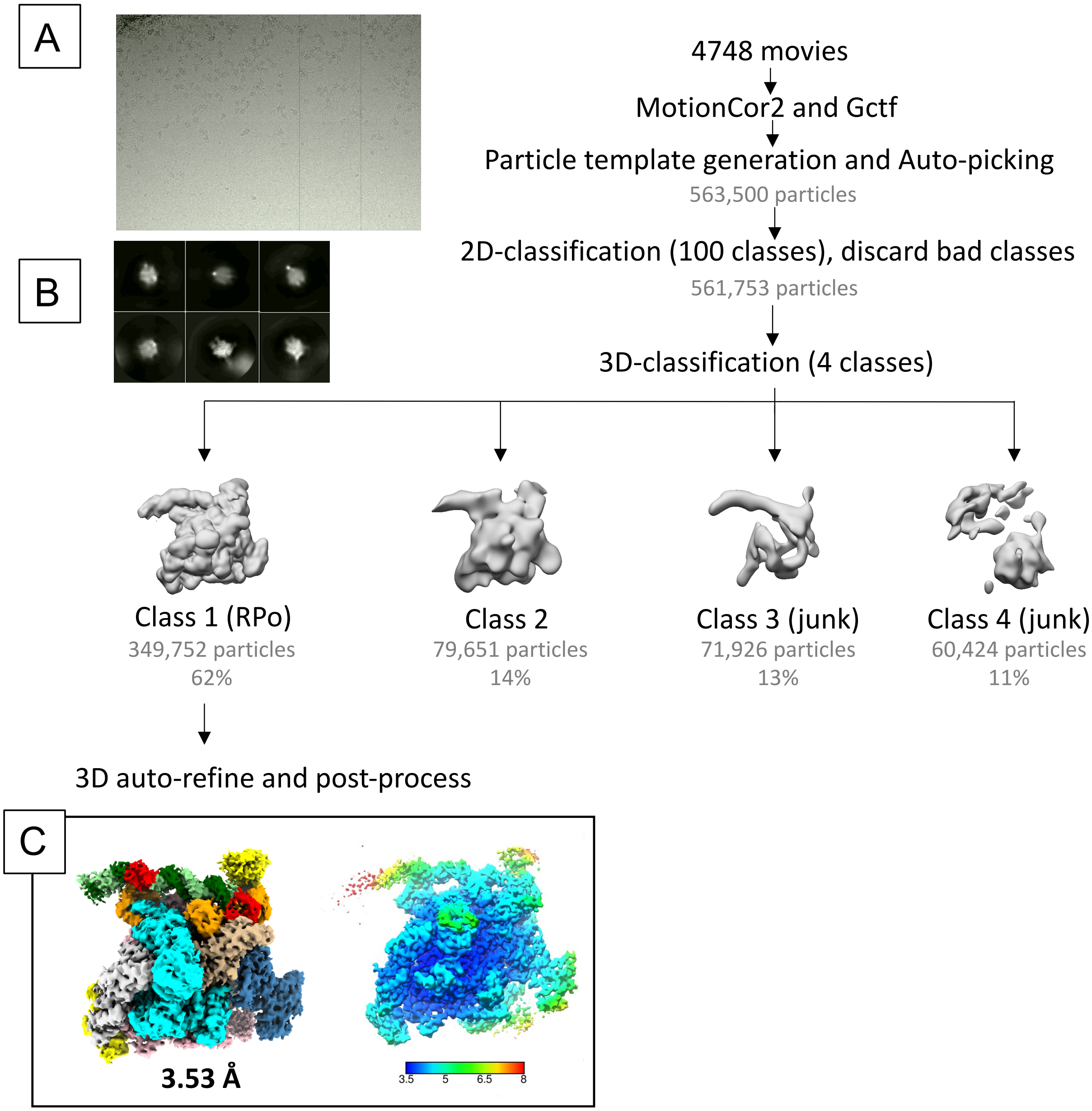

**SFigure 3. Cryo-EM processing pipeline for the RNAP-*rrnB*P1 complex.**

**A**) A representative micrograph used for data processing.

**B**) Selected representative 2D classes from 2D classification.

**C**) RPo. The cryo-EM density map is colored according to Fig. 2A. The right view is the same as on the left but colored by local resolution.

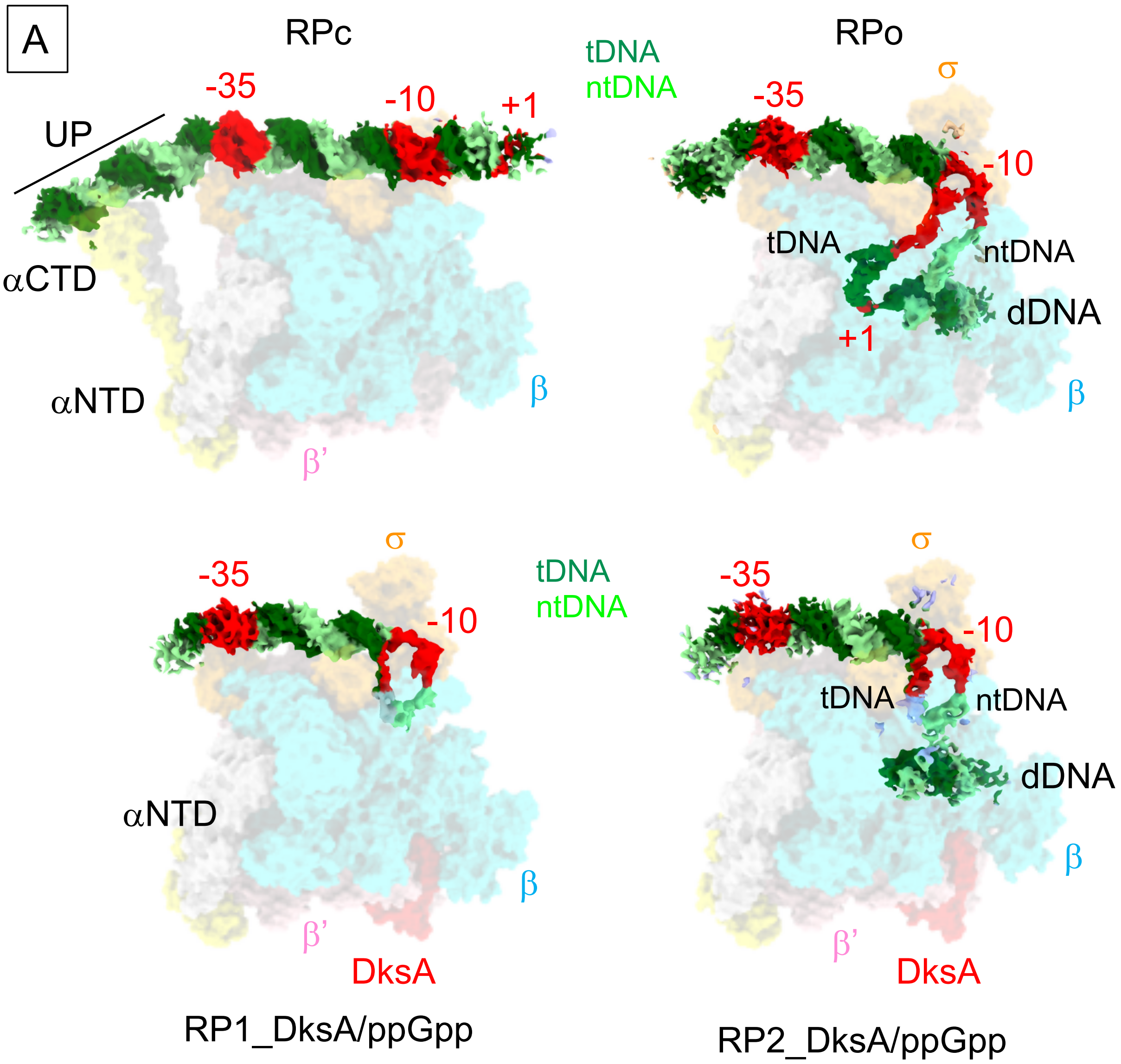

**SFigure 4. Cryo-EM density of promoter DNA from each structure.** (**A-D**) Cryo-EM density maps of the promoter DNA (tDNA, dark green; ntDNA, light green, -35 and -10 elements and +1 transcription start site, red) are shown with transparent RNAP density maps (α, yellow and white; β, cyan; β', pink; σ, orange; DksA, red). Active site Mg ions are shown as purple spheres. **A**) RPc. **B**) RPo. **C**) RP1-DksA/ppGpp. The cryo-EM density map is colored according to Fig. 4A. The right view is the same as on the left but colored by local resolution. **D**) RP2-DksA/ppGpp. The cryo-EM density map is colored according to Fig. 4B. The right view is the same as on the left but colored by local resolution.

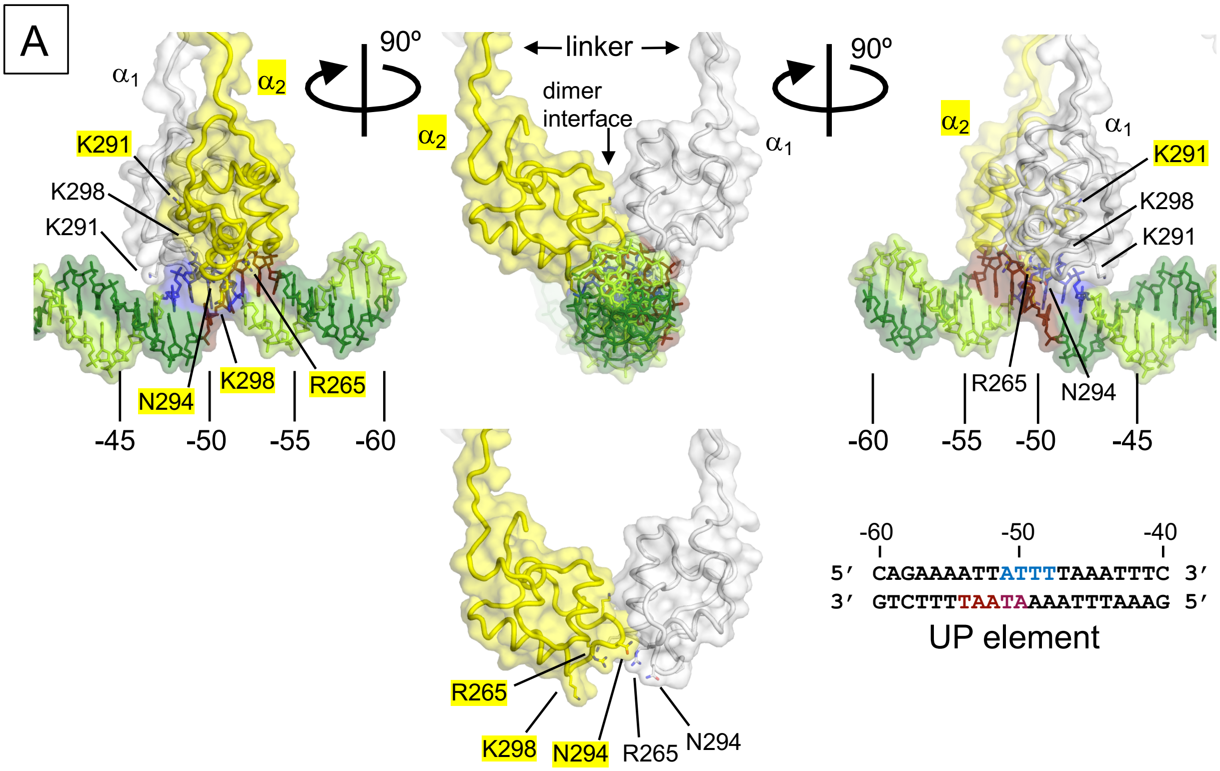

**SFigure 5. Detailed view of αCTDs-UP element interaction. Related to Fig. 1C.** Orthogonal views of RNAP contacting the UP element using the head-to-tail αCTD dimer. The αCTDs (yellow and white) and DNA around the UP element (dark and light greens) are depicted as cartoon and stick models, respectively. The amino acid residues of the αCTDs and the middle of the UP element (-51 to -48 on ntDNA, blue; -54 to -50 on tDNA, brown) making the interaction are indicated at the bottom.

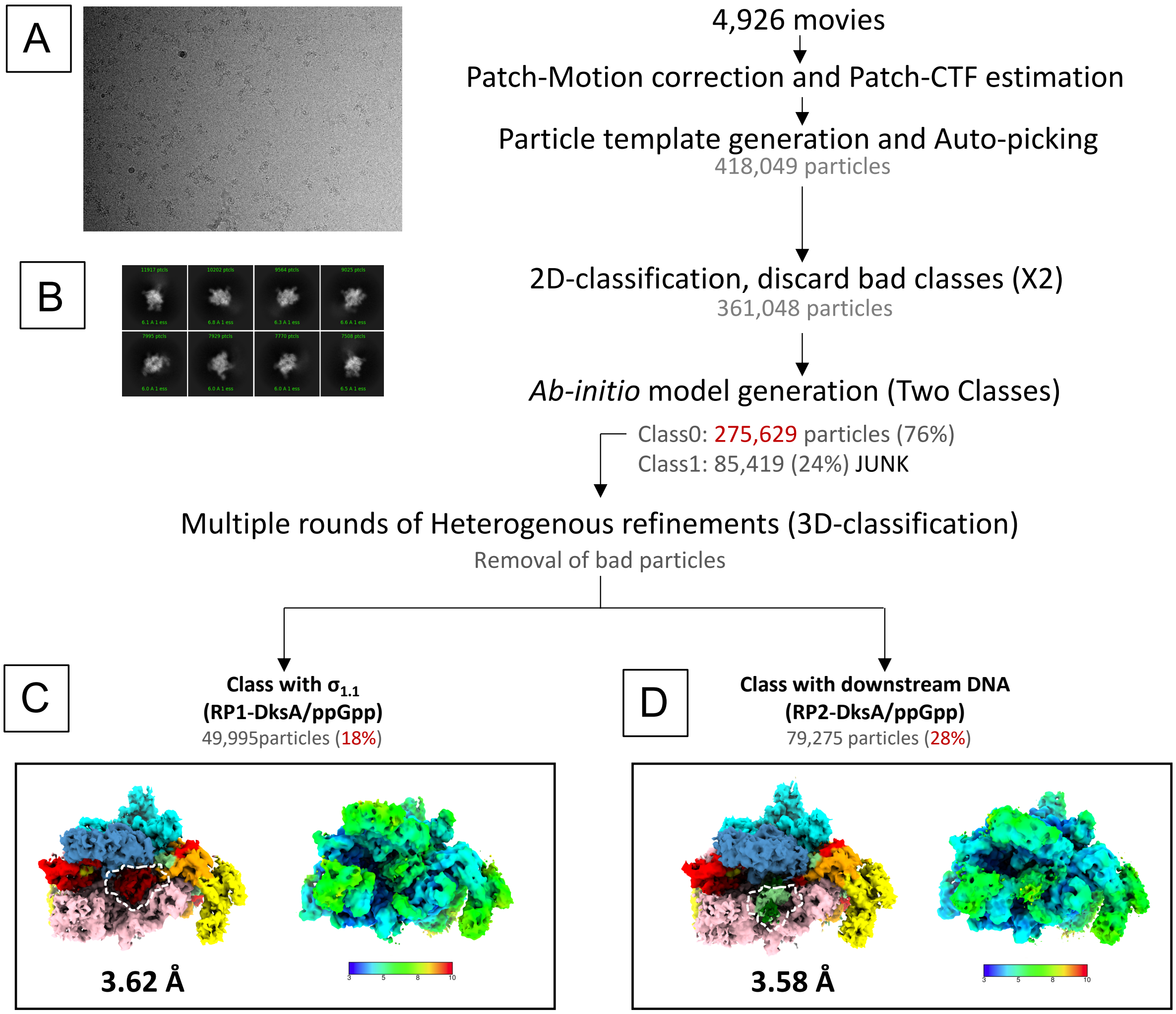

**SFigure 6. Cryo-EM data processing pipeline for the RP-DksA/ppGpp complex. A**) A representative micrograph of the RP-DksA/ppGpp complex used for data processing. **B**) Selected representative 2D classes from 2D classification. **C**) RP1-DksA/ppGpp, related to Fig. 4. **D**) RP2-DksA/ppGpp, related to Fig. 4.

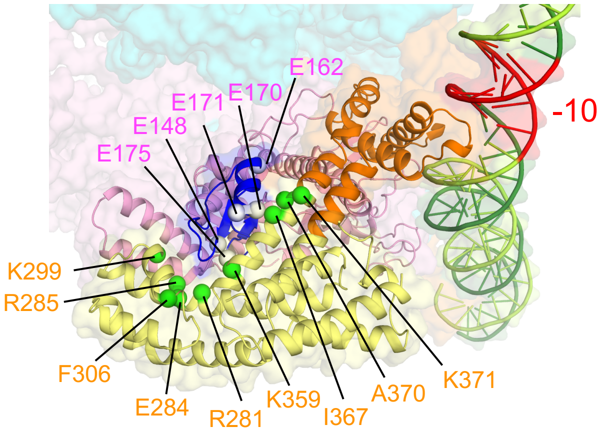

**SFigure 7. Magnified view of the $\sigma_{NCR}$ and β'clamp-toe (β'CT) interaction in the RPc structure.** The amino acid residues predicted to be involved in the $\sigma_{NCR}$ - β'CT interaction from the previous biochemical study [4] are shown as spheres and labeled.

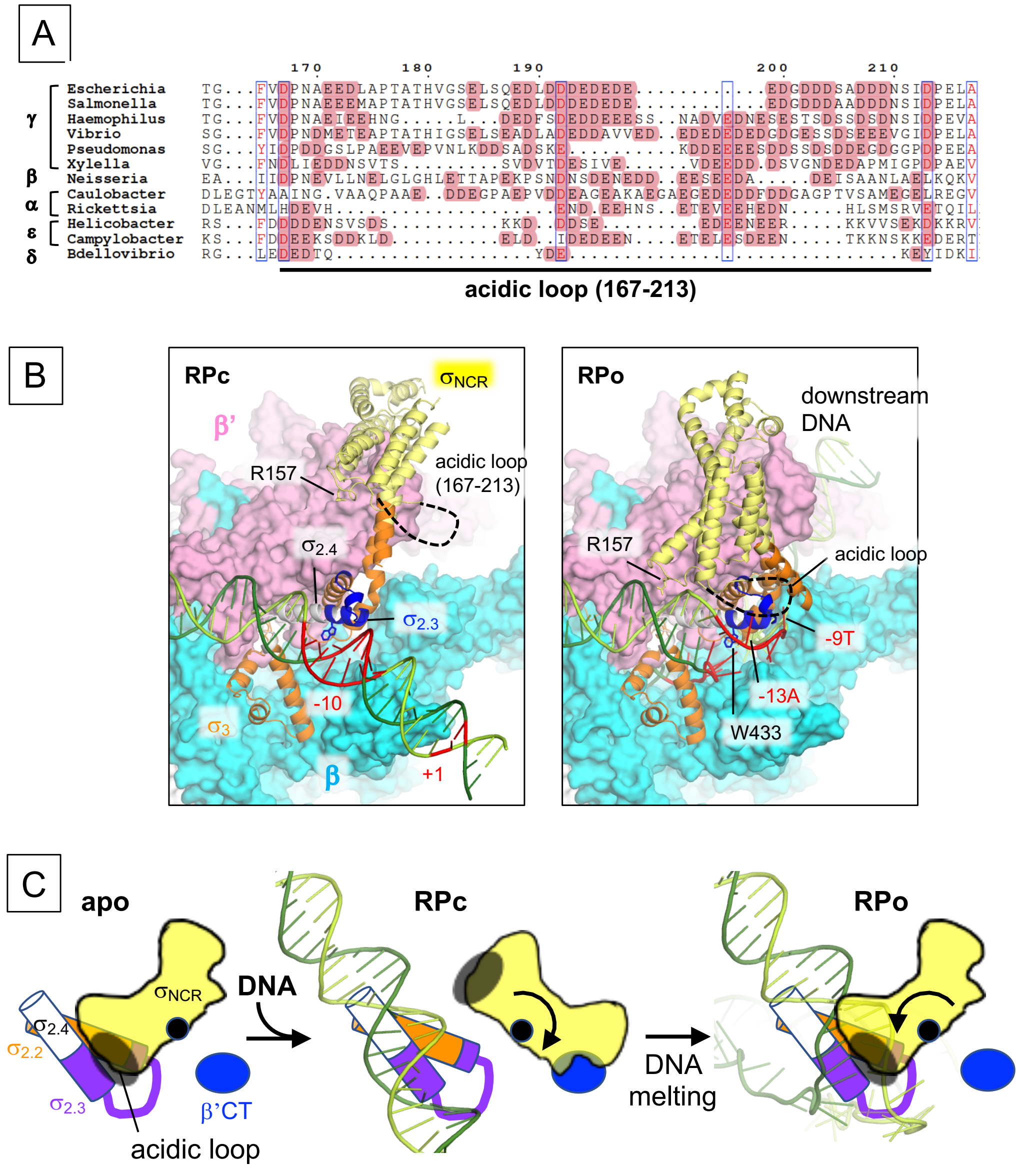

**SFigure 8. Sequence, structure and function of the acidic loop of $\sigma_{NCR}$. A**) Alignment of the $\sigma_{NCR}$ region shows the conservation of the acidic loop in proteobacteria. Acidic residues are highlighted in magenta. **B**) 3D structures showing the σ and -10 element interactions in RPc and RPo. The acidic loop is indicated as a black dashed line. **C**) A proposed role of the acidic loop of $\sigma_{NCR}$ in DNA binding and opening. $\sigma_{NCR}$ (yellow), σ domain 2 ($\sigma_{2.2}$, orange; $\sigma_{2.3}$, purple; $\sigma_{2.4}$, white), β'CT (blue) and DNA (light and dark green) are shown as a cartoon model. In the apo-form RNAP, the acidic loop (black) of $\sigma_{NCR}$ masks $\sigma_{2.3}$, which is unmasked upon the $\sigma_{NCR}$ - β'CT interaction during RPc formation. After DNA binding, the $\sigma_{NCR}$ interaction with DNA may facilitate DNA unwinding.

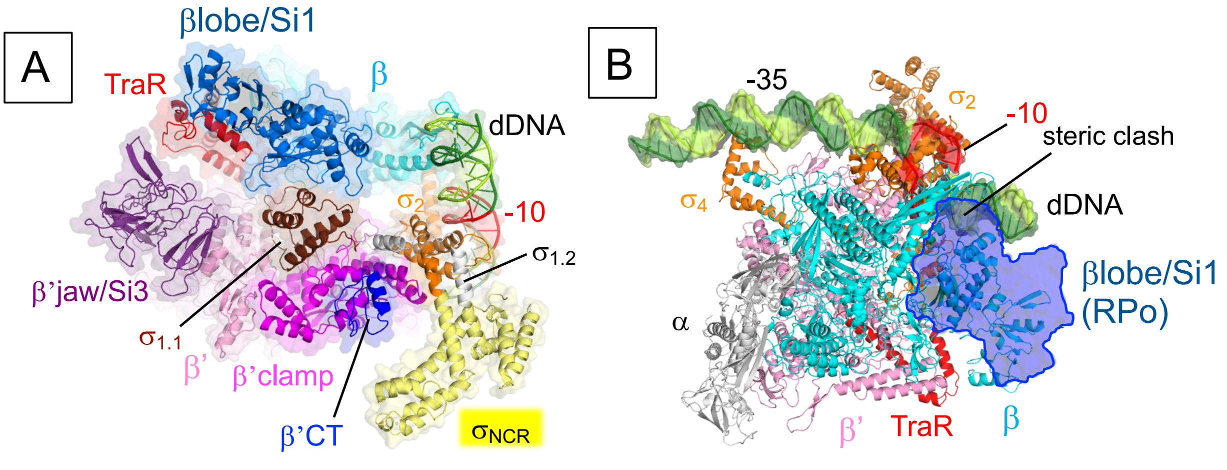

TRPc (PDB: 6PSQ)

TRPi1.5b (PDB: 6PST)

**SFigure 9. Cryo-EM structures of the RNAP – *rpsT*P2 complex with TraR in a closed complex form (TRPc) and an intermediate (TRPi1.5b)** [5]**.** RNAP (subunits and domains) and the *rpsT*P2 promoter DNA are shown as cartoon models with transparent surfaces. **A**) The structure of TRPc (PDB: 6PSQ) showed that $\sigma_{1.1}$ and $\sigma_{NCR}$ maintain their positions as in apo-form RNAP. **B**) The structure of TRPi1.5b (PDB: 6PST) highlighting a discriminator DNA accommodation above the βlobe/Si1 domain (cyan) in the presence of TraR. The conformation of the βlobe/Si1 domain in the absence of TraR (outlined in blue, RPo) precludes the discriminator DNA loading above the βlobe/Si1 domain.

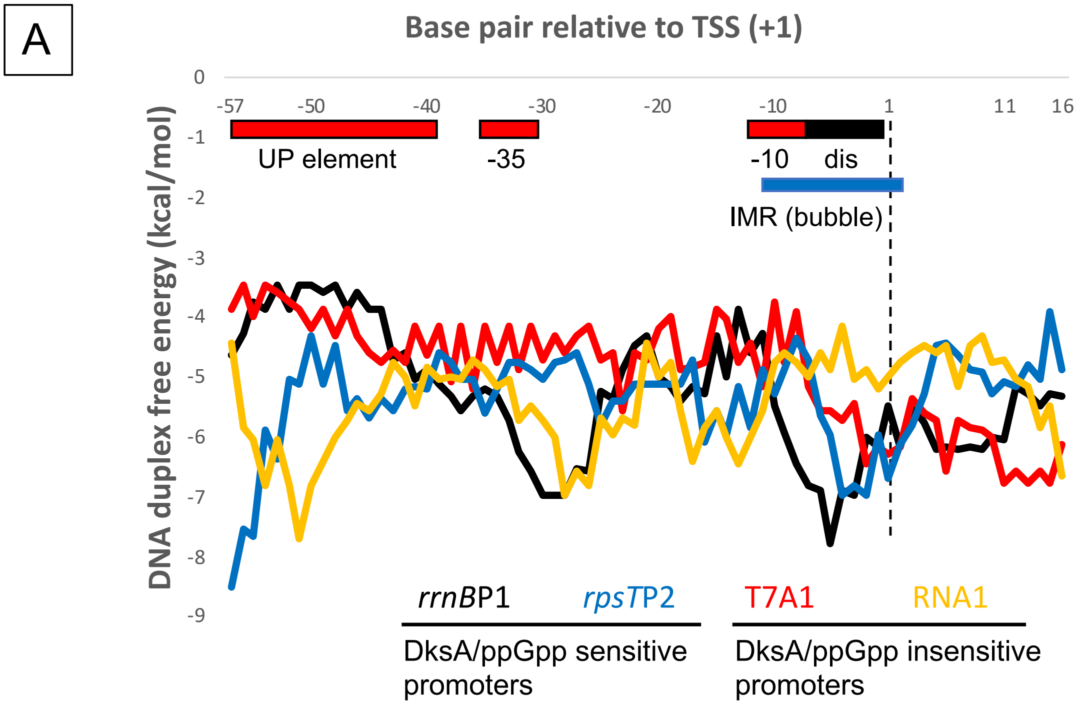

B

```
                                    discriminator
                    -35              -10       +1
5’ CAGAAAATTATTTTAAATTTCCTCTTGTCAGGCCGGAATAACTCCCTATAATGCGCCACCACTGACACGGACTCTACGAG 3’ rrnB P1
3’ GTCTTTTAATAAAATTTAAAGGAGAACAGTCCGGCCTTATTGAGGGATATTACGCGGTGGTGACTGTGCCTGAGATGCTC 5’

5’ GGCGGCGCTTATTTGCACAAATCCATTGACAAAAGAAGGCTAAAAGGGCATATTCCTCGGCCTTTGAATTGTCCATATAG 3’ rpsT P2
3' CCGCCGCGAATAAACGTGTTTAGGTAACTGTTTTCTTCCGATTTTCCCGTATAAGGAGCCGGAAACTTAACAGGTATATC 5’

5’ ATTTAAAATTTATCAAAAAGAGTATTGACTTAAAGTCTAACCTATAGGATACTTACAGCCATCGAGAGGGACACGGGGAA 3’ T7A1
3' TAAATTTTAAATAGTTTTTCTCATAACTGAATTTCAGATTGGATATCCTATGAATGTCGGTAGCTCTCCCTGTGCCCCTT 5’

5’ TATGTAGGCGGTGCTACAGAGTTCTTGAAGTGGTGGCCTAACTACGGCTACACTAGAAGAACTGTATTTGGTATCTGCGC 3’ RNA1
3’ ATACATCCGCCACGATGTCTCAAGAACTTCACCACCGGATTGATGCCGATGTGATCTTCTTGACATAAACCATAGACGCG 5’
```

**SFigure 10. DNA duplex free energy. A**) DNA duplex free energy calculated based on nearest-neighbor thermodynamics [6] for *rrnB*P1 (black), *rpsT*P2 (blue), T7A1 (red) and RNA-I (yellow) promoters. DNA sequences are shown in (**B**). Sequences were aligned at the promoter’s TSS. The UP element, -35 and -10 elements, discriminator (dis), and initially melted region (IMR or transcription bubble) are indicated.

**SMovie 1.** Cryo-EM density map of the RNAP - *rrnB*P1 closed complex (RPc). Related to Fig. 1B.

**SMovie 2.** Cryo-EM density maps of the RNAP - *rrnB*P1 open complex (RPo) and *rrnB*P1 DNA. Related to Fig. 2A.

**SMovie 3.** Close-up views of RNAP and discriminator DNA interactions. Related to Figs. 2C and D.

**SMovie 4.** Cryo-EM structures of the RNAP - *rrnB*P1 complex with DksA/ppGpp (RP1-DksA/ppGpp and RP2-DksA/ppGpp). Close-up view of the βlobe/Si1 conformational changes upon DksA binding, $\sigma_{1.1}$ ejection and downstream DNA binding. Related to Fig. 4.

**SMovie 5.** Alternative pathways for open promoter complex formation. Related to Fig. 6.

#### EXTENDED DATA REFERENCES

1 Condon, C., Philips, J., Fu, Z. Y., Squires, C. & Squires, C. L. Comparison of the expression of the seven ribosomal RNA operons in Escherichia coli. *EMBO J* **11**, 4175-4185, (1992).

2 Kolmsee, T., Delic, D., Agyenim, T., Calles, C. & Wagner, R. Differential stringent control of Escherichia coli rRNA promoters: effects of ppGpp, DksA and the initiating nucleotides. *Microbiology* **157**, 2871-2879, (2011).

3 Aiyar, S. E., Gaal, T. & Gourse, R. L. rRNA promoter activity in the fast-growing bacterium Vibrio natriegens. *J Bacteriol* **184**, 1349-1358, (2002).

4 Leibman, M. & Hochschild, A. A sigma-core interaction of the RNA polymerase holoenzyme that enhances promoter escape. *EMBO J* **26**, 1579-1590, (2007).

5 Chen, J. *et al.* Stepwise Promoter Melting by Bacterial RNA Polymerase. *Mol Cell*, (2020).

6 SantaLucia, J., Jr. A unified view of polymer, dumbbell, and oligonucleotide DNA nearest-neighbor thermodynamics. *Proc Natl Acad Sci U S A* **95**, 1460-1465, (1998).
